# Nonlinear mixed selectivity supports reliable neural computation

**DOI:** 10.1101/577288

**Authors:** W. Jeffrey Johnston, Stephanie E. Palmer, David J. Freedman

**Affiliations:** Graduate Program in Computational Neuroscience, The University of Chicago, Chicago, IL, USA; Department of Neurobiology, The University of Chicago, Chicago, IL, USA; Department of Organismal Biology and Anatomy, The University of Chicago, Chicago, IL, USA; Department of Physics, The University of Chicago, Chicago, IL, USA

**Author notes:** Equal contributions (Stephanie E. Palmer, David J. Freedman).

**Keywords:** neural coding, reliability, efficiency, nonlinear mixed selectivity, conjunctive coding

## Abstract

Neuronal activity in the brain is variable, yet both perception and behavior are generally reliable. How does the brain achieve this? Here, we show that the conjunctive coding of multiple stimulus features, commonly known as nonlinear mixed selectivity, may be used by the brain to support reliable information transmission using unreliable neurons. Nonlinear mixed selectivity (NMS) has been observed widely across the brain, from primary sensory to decision-making to motor areas. Representations of stimulus features are nearly always mixed together, rather than represented separately or with only additive (linear) mixing, as in pure selectivity. NMS has been previously shown to support flexible linear decoding for complex behavioral tasks. Here, we show that NMS has another important benefit: it requires as little as half the metabolic energy required by pure selectivity to achieve the same level of transmission reliability. This benefit holds for sensory, motor, and more abstract, cognitive representations. Further, we show experimental evidence that NMS exists in the brain even when it does not enable behaviorally useful linear decoding. This suggests that NMS may be a general coding scheme exploited by the brain for reliable and efficient neural computation.

## Introduction

To support behavior, the brain must use a communication strategy that transmits information about the world faithfully, efficiently, and, perhaps most of all, reliably. The first two of these goals have received extensive attention in neuroscience, particularly in the literature on efficient coding and redundancy reduction[1]. Efficient coding focuses on discovering the response field (RF) for a single neuron that simultaneously maximizes the amount of stimulus information transmitted by the neuron while minimizing the number of spikes that the neuron must fire[1]. A crucial step to this process is representing stimuli without any of the redundancy inherent to the natural world – that is, by isolating and representing the independent components of natural stimuli[2]. Refinements of efficient coding[3] have also emphasized the need for the representation of these components to be neatly packaged, or formatted, so that they are accessible to decoding (as with nonlinear mixed selectivity[4]) and facilitate generalization[5]. As a whole, the ideas of efficient coding have been used to accurately predict the structure of RFs in primary visual cortex[6, 7], and other sensory systems[8–10]. However, efficient coding, redundancy reduction, and neat packaging all address only the goals of faithful representation and metabolic efficiency. They do not guarantee that these transmissions will be reliable when corrupted by the noise that is inherent to single neuron responses[11, 12]. In fact, non-redundant representations are often highly vulnerable to noise[13].

Making efficient representations robust to the noise present throughout the neural system – and satisfying the final goal, transmission reliability – has received considerably less attention in neuroscience. In information theory, noise robustness is the goal of channel coding, which re-codes efficient stimulus representations to include redundancy that is specifically useful for improving transmission reliability. Recent work has shown that grid cell RFs[14] and the working memory system[15] may implement near-optimal channel codes. In sensory systems, channel coding has been explored more obliquely. Extensive work has focused on deriving RF properties that maximize mutual information between the stimulus and the response[16] or the Fisher information from the response function[17–19] (and see [20, 21] for connections between these approaches). However, neither of these measures has a direct connection to decoder performance: mutual information is linked via the rate-distortion bound[22], but high mutual information does not guarantee good decoder performance in general[23] (and see Figure 2C); Fisher information is linked via the Cramer-Rao bound, but saturation of this bound is only guaranteed in low-noise conditions[24] (and codes with less Fisher information can outperform codes with more Fisher information when optimal decoding cannot achieve the bound[25]). There is neural and behavioral[26] evidence that the brain computes successfully on short (e.g., ~80 ms) timescales and spiking responses have been shown to be highly variable on that timescale[26], thus it is unlikely that the brain typically operates in a low-noise regime.

Here, we analyze an ubiquitous coding strategy in the brain – conjunctive coding for multiple stimulus features – in terms of both its reliability and efficiency. Previous work on conjunctive coding (commonly called nonlinear mixed selectivity[4, 27]) has shown that it produces a neatly packaged and sparse representation that enables the use of simple linear decoders for complex cognitive tasks[4], particularly in the macaque prefrontal cortex[27]. Further, random conjunctive coding has been shown to increase the number of discrete stimuli that can be reliably represented in a neural population[28, 29]; however, a detailed analysis of how the error rate of these codes depends on metabolic cost was not performed. In our work, we develop a novel generalization of nonlinear mixed selectivity (NMS), allowing different levels of mixing between stimulus features while preserving full coverage of the stimulus space (see *Definition of the codes* in *Methods*). Using these codes, we show that the encoding of stimuli with at least some level of NMS almost always produces more reliable and efficient communication than without NMS. Further, we demonstrate novel tradeoffs between codes with and without NMS – including an analysis of how RF size and error-type affect the optimal level of NMS. Finally, we link our work to experimental data by showing that NMS is implemented in the brain when it could support reliable readout without playing a role in the linear decoding of stimulus features. Our work illustrates that NMS provides highly general benefits to coding reliability and efficiency, and helps to explain the ubiquity of NMS within sensory[30–35], frontal[4, 27], and motor cortices[36–38].

## Results

### Increased mixing increases stimulus discriminability

In the brain, stimulus representations are corrupted by noise as they are transmitted between different neural populations. This process can be formalized as transmission down a noisy channel (Figure 1A). The reliability and efficiency of these transmissions depends on the format of the encoded representations – here, we show how three different properties of this representation are affected by NMS, and how those properties interact with transmission reliability and efficiency. The three properties of neural representations that we focus on are: minimum distance, neural population size, and metabolic representation energy (Figure 1B and *Code properties* in *Methods*). Minimum distance is the distance between representations of two stimuli that are most difficult to discriminate. Importantly, half the minimum distance represents the smallest magnitude of noise that could cause a decoding error given optimal decoding. A larger minimum distance typically implies a lower overall probability of error (see *Union bound estimate* in *Methods*), even for non-optimal decoding. Population size is the minimum number of independent coding units, or neurons, required to implement the code such that all possible stimuli have a unique response pattern. Representation energy is the metabolic energy consumed by a stimulus response in the code, defined as the square of the distance between the zero-activity state and all the response patterns in the code – here, representation energy can be viewed as the squared spike rate in response to a particular stimulus summed across the population of neurons used by the code (though we also consider the sum spike rate, see Figure S1). In the codes we consider here, all of our stimuli evoke the same summed spike rate across the population, and therefore have the same representation energy. Representation energy represents the active, metabolic cost of the code (in terms of the cost of emitting spikes), while population size represents the passive metabolic cost of the code (in terms of neuronal maintenance costs across the population, spiking or not). We begin by considering representation energy alone before considering both together.

**Figure 1:**
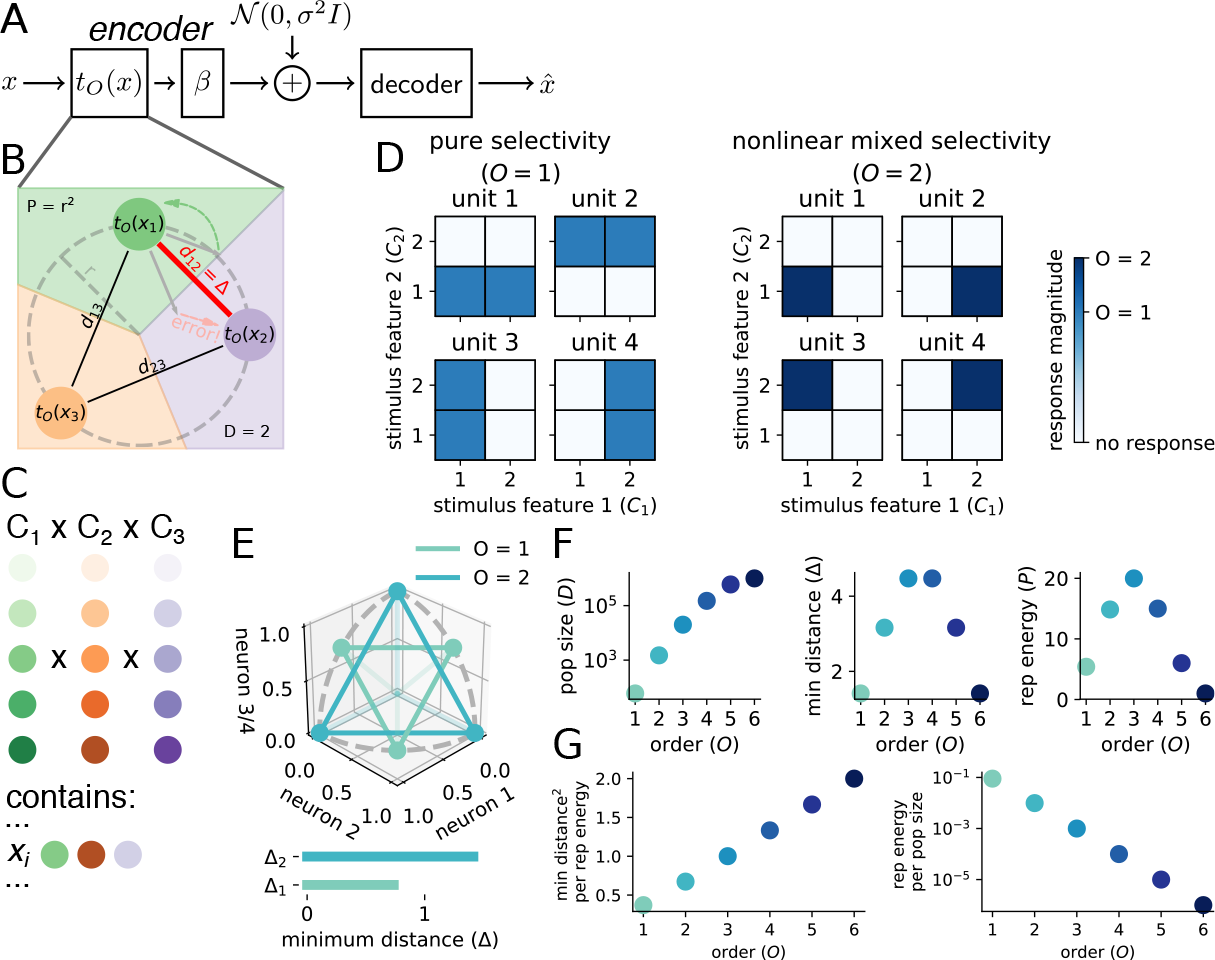
Mixed codes produce more discriminable stimulus representations. **A** The noisy channel model. A stimulus *x* is encoded by encoding function *t_O_* (*x*) of order *O* then amplified by linear transform *β* before independent Gaussian-distributed noise with variance *σ*^2^ is added in the channel. Next, a decoder produces an estimate 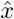 of the original stimulus. **B** We analyze the encoding function with respect to three important code properties. The minimum distance Δ = *d*_12_ is the smallest distance between any pair of encoded stimuli (codewords), and half of that distance is the nearest border of the Voronoi diagram (background shading). Thus, minimum distance can be used to approximate the probability of decoding error. Representation energy *P* = *r*^2^ is the square of the radius of the circle that all of the codewords lie on. All of the codewords lie in a 2-dimensional plane, so the code has population size *D* = 2. **C** Stimuli are described by *K* features *C_i_* which each take on *C*_*i*_ = *n* values. All possible combinations of feature values exist, so there are *n^K^* unique stimuli. **D** In pure selectivity (left), units in the code, or neurons, respond to a particular value of one feature and are invariant to changes in other features. In nonlinear mixed selectivity (right), neurons respond to particular combinations of feature values, and the number of feature values in those combinations is defined as the order *O* of the code (here, *O* = 2). **E** The same *O* = 1 and *O* = 2 code as in **D**. (top) The colored points are the response patterns in 3D response space for three of the four neurons in each code. The dashed grey line is the radius of the unit circle centered on the origin for each plane – the two codes are given constant representation energy, and all response patterns lie on the unit 4D hypersphere. For ease of visualization, the vertical dimension in the plot represents both the third and fourth neurons in the population to show three representations from the *O* = 1 code, this does not change the minimum distance. (bottom) The response patterns for the *O* = 2 mixed code have greater minimum distance than those for the *O* = 1 pure code. **F** We derive closed-form expressions for each code metric, and plots of each metric are shown for codes of order 1 to 6 with *K* = 6 and *n* = 10. **G** Mixed codes produce a higher minimum distance per unit representation energy (left) and have a smaller proportion of active units (i.e., greater sparseness; right) than pure codes.

The stimuli represented by our codes are described by *K* independent features that each take one of *n* discrete values with equal probability (Figure 1C and see *Definition of the stimuli* in *Methods*). As a simple example, one feature could be shape, and two values for shape could be square or triangle; a second feature could be color, and two values could be red or blue. In all, there are *n^K^* possible stimuli. So, four stimuli in our example. Each of the stimuli are equally likely. While we focus on discrete features, our core result is the same with continuous features (see *Error-reduction by NMS in the continuous case* in *Supplemental Information*).

To understand how NMS affects code reliability and efficiency, we compare the performance of codes with different levels of conjunctive stimulus feature mixing (i.e., different code orders; Figure 1D), following the definition of NMS used previously in the literature[4, 27]. Neurons in a code of order *O* respond to a particular combination of *O* feature values and do not respond otherwise (Figure 1D and see *Definition of the codes* in *Methods*), and a code has a neuron that responds to each possible combination of *O* different stimulus feature values (see *Code example* in *Methods* for more details). In our example, an order one (*O* = 1) code would have neurons that respond to each shape regardless of color and each color regardless of shape while an order two code (*O* = 2) would have neurons that respond to each combination of shape and color – for instance, one neuron would respond only to red squares, another only to blue squares, and so on. This example can map onto the two features used in the illustration in Figure 1D, E. From this construction, each stimulus will have a unique response pattern across the population of neurons, but the population size will vary across code order. In general, higher-order codes will have larger population sizes, but sparser neural responses (i.e., a smaller percentage of neurons will respond to each stimulus).

To test this intuition in general, we derive closed-form expressions for the minimum distance (Δ_*O*_), population size (*D_O_*), and representation energy (*P_O_*) of our codes that depend on the number of features *K*, the number of values each of those features can take on *n*, and the order of the code *O* (Figure 1F and *Code properties* in *Methods*). Using these expressions, we show that the ratio between squared minimum distance Δ_*O*_ and representation energy *P_O_* is strictly increasing with order for all choices of *K* and *n* (see *Minimum distance-representation energy ratio* in *Methods* and Figure 1G, left):

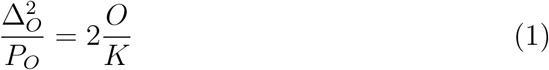

This shows that, given the same amount of representation energy, codes with more mixing produce stimulus encodings with strictly larger minimum separation in the response space. This is suggested by their increased sparseness (Figure 1G, right). While sparseness per se does not have a direct relationship with code performance, increased sparseness indicates that the response patterns used by an encoding scheme may (but do not necessarily) have less overlap and therefore greater distance from each other in response space than a denser representation. In our codes, the increase in sparseness produced by an increase in code order does produce greater response separation (Figure 1E illustrates this effect). This increased separation provides a benefit to decoding for many different noise distributions and decoders (including linear and maximum likelihood decoders), and indicates that mixed codes are likely to produce more reliable and efficient representations than pure codes in a wide variety of conditions. However, to directly quantify transmission reliability (i.e., the probability of a decoding error), we must include the details of both the noise and the decoder (see Figure 1A and *Full channel details in Methods*).

### Mixed codes make fewer errors than pure codes

To directly estimate the probability of decoding error (PE) for each of our codes, we expand our analysis from the encoding function (Figure 1) to the channel as a whole. We choose the noise to be additive, independent, and Gaussian (though we also consider multiplicative, Poisson noise, which gives similar results, see Figure S2) and use a maximum likelihood decoder (MLD, and see *Full channel details* in *Methods*). Given these noise and decoder assumptions, we can use the union bound estimate (UBE) to approximate PE (UBE) of PE (see *Union bound estimate* in *Methods*). The UBE decomposes the probability that we make an error into the sum of the probabilities of only the most likely errors (that is, errors to stimuli at minimum distance). This estimate takes the form:

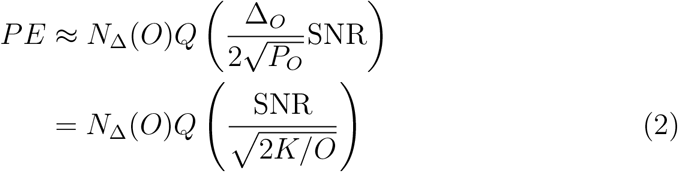

where *Q*(*y*) is the complementary cumulative distribution function of 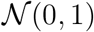 at *y*, 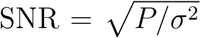 is the signal-to-noise ratio (see *The amplifying linear transform (β)* in *Methods*), and *N*_Δ_(*O*) is the number of neighbors at minimum distance of the order *O* code, which is equal to

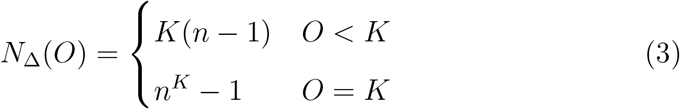

see *Code neighbors* in *Supplemental Information*. Thus, for constant SNR, our estimate of the probability of decoding error is strictly decreasing with order for *O* < *K*, but not necessarily for *O* = *K* when *n^K^* is large.

To verify that the UBE provides a good estimate of PE, we numerically simulate codes of all possible orders over a wide range of SNRs for particular choices of *K* and *n* using the same channel as in our analysis. Our simulations show that higher-order codes outperform lower-order codes across all SNRs at which the codes are not saturated at chance or at zero error (Figure 2A). We also show that the UBE closely follows performance for large SNRs (Figure 2A inset). Using the UBE, we compare the amount of representation energy that codes of different orders require to reach 1% decoding error (Figure 2B) and show that a pure code requires up to twice as much representation energy as a mixed code with the optimal order. This also illustrates that while the nearest-neighbor increase for the full-order code is not a practical concern for low *n^K^* (as the number of neighbors is still small), it quickly causes the *O* = *K* − 1 code to outperform the *O* = *K* code as *n^K^* becomes large (Figure 2B). In all conditions we simulated (following the UBE), either the *O* = *K* − 1 or *O* = *K* code provided the lowest decoding error at a given SNR. Thus, in these conditions, mixed codes provide a significant benefit to coding reliability independent of particular parameter choices.

**Figure 2:**
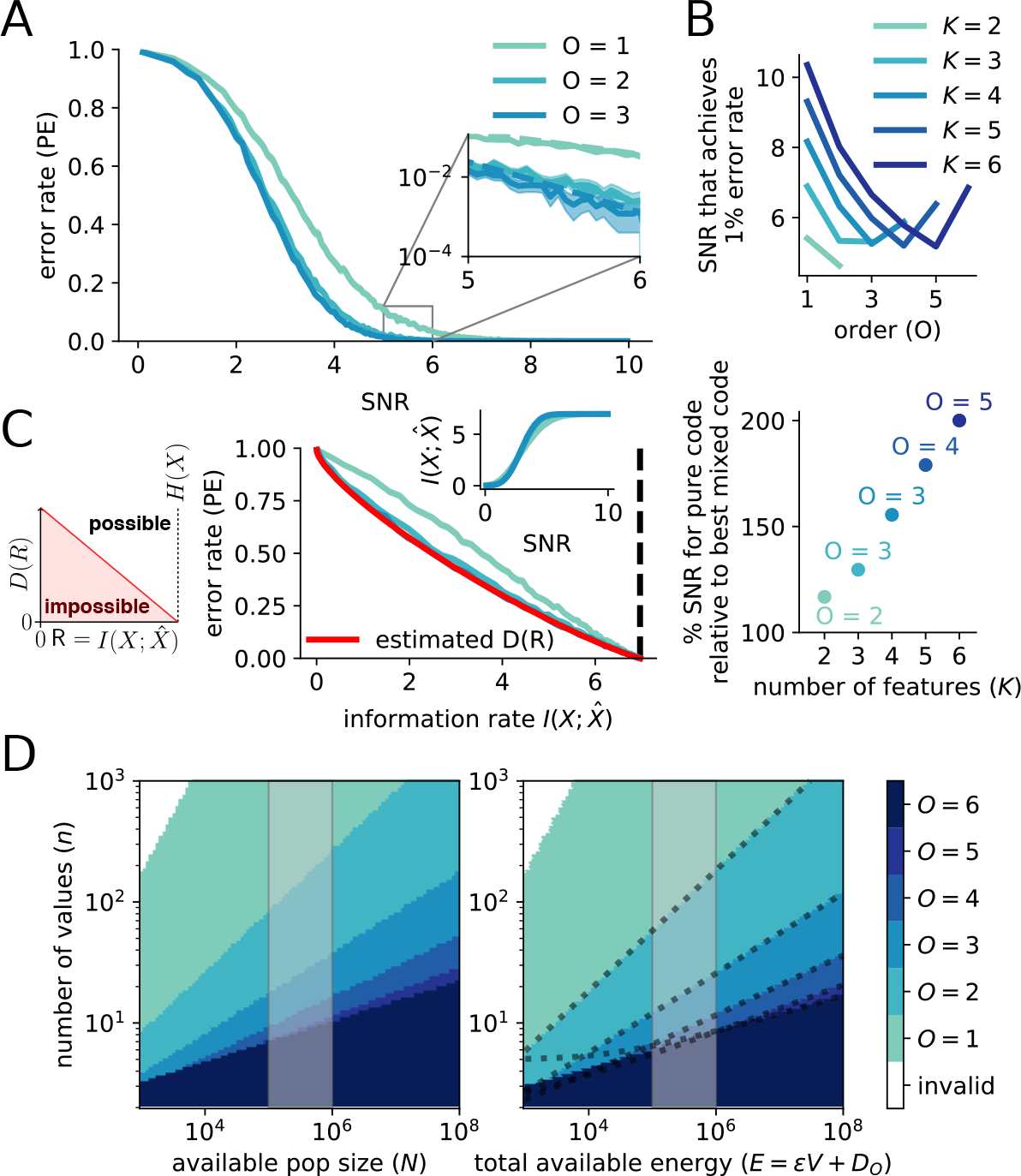
Mixed codes make fewer errors than pure codes. **A** Simulation of codes with *O* = 1, 2, 3 for *K* = 3 and *n* = 5. (inset) For high SNR, code performance is well-approximated by the union bound estimate (UBE). **B** (top) Using the UBE, we show that for different *K* (with *n* = 5) the SNR required to reach 1 % decoding error tends to decrease with increasing code order. (bottom) The representation energy required by the pure code relative to that required by the best mixed code (given by point color and label) to reach 1 % decoding error. **C** (left) A schematic of the distortion-rate function for some source distribution *X*. Through rate-distortion theory, we know that no codes below the distortion-rate bound will ever be discoverable for this particular form of source distribution *X*. (right) With *K* = 3 and *n* = 5, the performance of different order codes relative to the bound. (right, inset) The codes have slightly different efficiencies for transforming SNR to 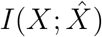 **D** (left) Given a pool of neurons with fixed size, the color corresponding to the code producing the highest minimum distance is shown in the heat map. The shaded area delineates the order of magnitude of the number of neurons believed to be contained in 1 mm^3^ of mouse cortex. (right) The same as on the left, but instead of a pool of neurons of fixed size, each code is given a fixed total amount of energy. The energy is allocated to both passive maintenance of a neural population (with size equal to the population size of the code) and representation energy (increasing SNR). The shaded area is the same as on the left. The dashed lines are plots of our analytical solution for the transition point between the *O* and *O* + 1-order code (see *Total energy* in *Methods*).

For smaller choices of *K* and *n*, we were able to empirically evaluate how decoding error compares to the rate-distortion bound[22]. In this context, the rate-distortion bound is an absolute lower bound on the probability of making a decoding error given a particular information rate through the channel (i.e., the mutual information 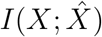 and see *The rate-distortion bound and mutual information calculation in Methods*). We first show that higher-order codes generate a higher information rate than lower-order codes at the same SNR (Figure 2C inset) – that is, they more efficiently transform the input into stimulus information. Next, we show that the full-order code (*O* = *K* = 3) fully saturates the rate-distortion bound (Figure 2C). Thus, for a given amount of stimulus information, full-order codes produce as few errors as would be possible for any code[22]. While the *O* = *K* − 1 = 2 code comes close to this bound as well, the pure code does not.

### Mixed codes provide benefits despite requiring more neurons

Our analysis so far has focused on the metabolic cost of neuronal spiking. A single spike is thought to be the largest individual metabolic cost in the brain[39]. For a fixed population size *N*, from Eq. 1, we know that the code with the highest order *O* such that *D*_*O*_ ≤ *N* will provide the largest minimum distance, given a fixed amount of spiking activity (Figure 2D). For a wide range of stimulus set sizes, mixed codes have population sizes less than or equal to an order-of-magnitude estimate of the neuron count in 1 mm^3^ of mouse cortex[40] (Figure 2D, shaded region). Thus, the benefits of mixed codes are practically achievable in the brain.

However, the passive maintenance of large neural populations also has a metabolic cost, due to the turnover of ion channels and other cell-level processes[39], which, for large populations of sparsely firing neurons, could be as large if not larger than the metabolic cost associated with spiking. To account for this cost, we adapt the formalization from [41] to relate representation energy (i.e., spiking) to the metabolic cost of population size (see *Total energy in Methods*). We refer to the sum of these costs as the total energy *E* of a code. Codes with small population sizes will be able to allocate more of their total energy to representation energy, while codes with large population sizes will have less remaining total energy to allocate to representation energy. We do not constrain the maximum SNR that a single neuron in our codes can achieve (even though achievable SNR is limited in the brain[42]), which further improves the performance of pure codes relative to higher-order codes under our formalization. Thus, it serves as a particularly stringent test of the reliability and efficiency of mixed codes.

Mixed codes yield higher minimum distance under the total energy constraint for a wide range of stimulus set sizes and total energy (Figure 2D), including order-of-magnitude estimates of the total energy available to 1 mm^3^ of mouse cortex (Figure 2D, shaded region). Further, our analysis reveals that for any total energy *E ≥ n*^2^*K*^2^ (see Eq. M.6) a mixed code (*O >* 1) will provide better performance than the pure code (*O* = 1). These results also make an important prediction that can be tested experimentally: the order of neuronal RFs should decrease as the fidelity required of the representation increases (i.e., as *n* increases). There already exists indirect experimental support for this prediction. In the visual system, single neurons in primary visual cortex have RFs thought to represent relatively small combinations (small *O*) of lowlevel stimulus features such as spatial frequency and orientation[6, 30] (but see [43]), while single neurons in the prefrontal cortex are thought to have responses that depend on larger combinations (high *O*) of abstract, often categorical (and therefore low *n*), stimulus features along with behavioral context[4, 27]. However, this pattern has not been rigorously tested, as these regions are rarely recorded in the same tasks and the tasks chosen for each area often follow the form of the prediction – that is, requiring high fidelity (*n*) for investigations of primary sensory areas and low fidelity (*n*) for investigations of prefrontal areas.

### Mixed codes provide reliable coding in sensory systems

So far, we have focused on the probability of decoding error, which is most applicable to features that represent categorical differences without defined distances from each other (e.g., mistaking a hat for a sock is not clearly less accurate than mistaking a hat for a glove). However, in sensory systems, the features often do have a relational structure and stimuli that are nearby to each other in feature space are also perceptually or semantically similar (e.g., mistaking a 90° orientation for a 180° orientation is clearly less accurate than mistaking 90° for 100°). In the context of sensory information, minimizing the frequency of errors becomes less important than ensuring that the average distance of an estimate from the original stimulus is low. This is because perceptually similar errors are likely more useful for guiding behavior than a random error, even if the latter occurs less frequently. This difference in priority is encapsulated in the contrast between PE (Figure 3A) and the mean squared-error distortion (MSE; Figure 3B), which is equivalent to the average squared-distance of the estimated stimulus from the original stimulus. In our framework, full-order mixed codes have the highest minimum distance (Eq. 1), but all stimuli are nearest neighbors to all other stimuli (Eq. 3) which causes all errors to be random with respect to the original stimulus. Thus, using MSE instead of PE, we show that lower-order mixed and pure codes outperform full-order mixed codes at low total energy (Figure 3B). However, increased total energy causes a faster decay in error rates for full-order codes than lower-order codes (Eq. 2). Thus, even full-order codes outperform pure codes under MSE at high total energy (Figure 3B).

**Figure 3:**
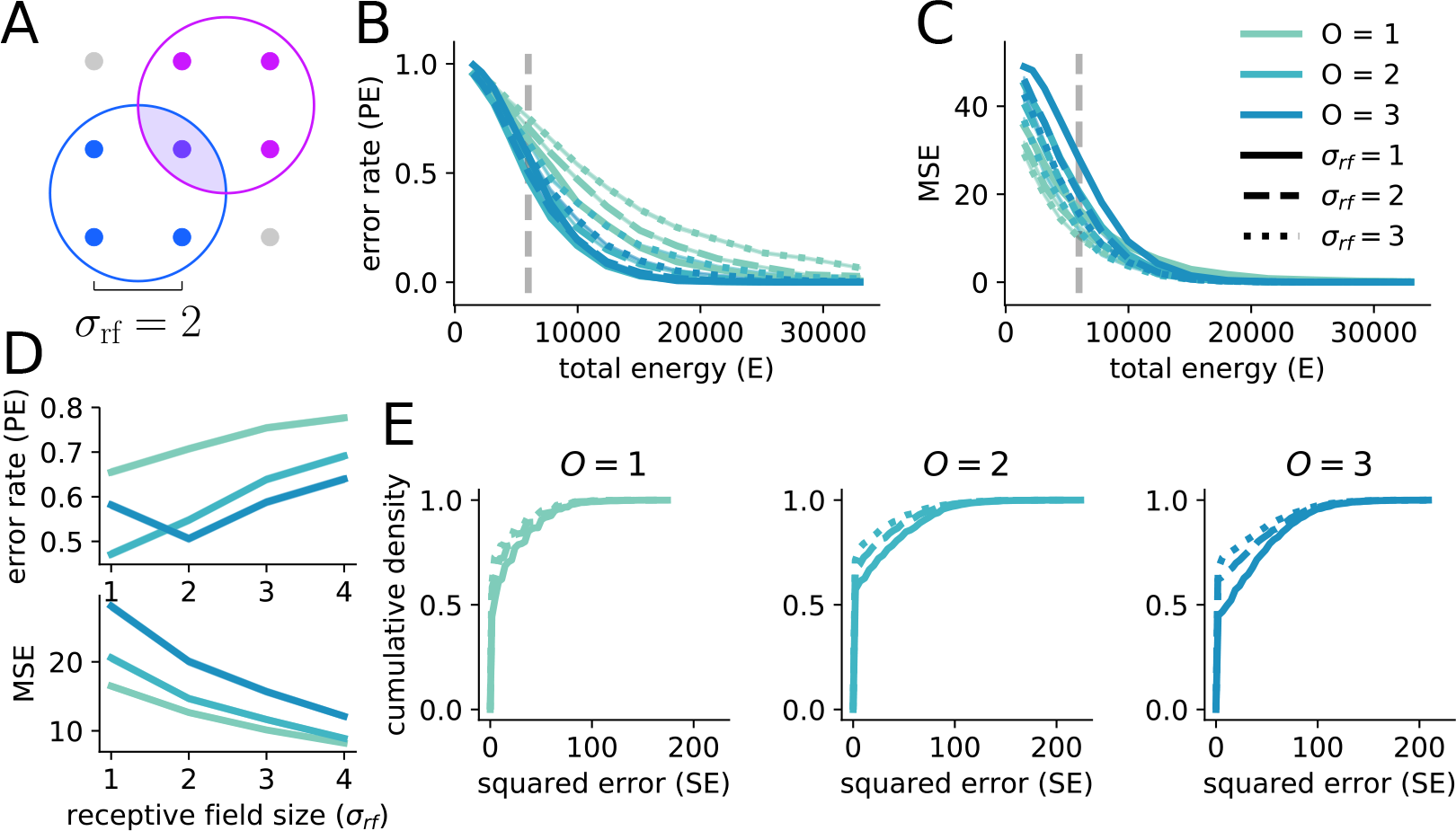
Mixed codes can be more reliable than pure codes for both PE and MSE, but different RF sizes are appropriate for each. **A** An illustration of our RF formalization. With *K* = 2 and *n* = 3, two example RFs of size *σ*_rf_ = 2 are shown. Simultaneous activity from both neurons uniquely specifies the center stimulus point. **B** Simulated PE of codes of all orders for *K* = 3 and *n* = 10 with *σ*_rf_ = 1, 2, 3 (legend as in **C**). Note that total energy is plotted on the x-axis, rather than SNR as in Figure 2. Mixed codes outperform the pure code over many (but not all) total energies. **C** The same as **B** but for MSE rather than PE. Mixed codes perform worse than pure codes for low total energy, but perform better as total energy increases. **D** PE increases (top) and MSE decreases (bottom) as *σ*_rf_ increases for the codes in **B** and **C** taken at the total energy denoted by the dashed grey line. **E** Cumulative distribution functions for the squared errors made for the codes given in **B** and **C** at the grey dotted line. MSE is decreased by increasing *σ*_rf_ despite the increase in PE because the errors that are made become smaller in magnitude and this outweighs their becoming more numerous. This effect is largest for the *O* = *K* = 3 code.

Further, we show that the brain can take advantage of the increased minimum distance yielded by full-order codes and reduce the randomness of errors in the context of sensory systems by increasing the size of neuronal RFs (Figure 3C), which also makes higher-order codes more practical by reducing their population size (see *Additional results on response fields* in *Supplemental Information*). Thus, this work provides a unified framework for understanding the purpose and benefits of large RFs in arbitrary feature spaces, which are often observed in cortex[44]. In particular, increasing RF size decreases the MSE for all codes while increasing the PE (Figure 3D and see *Additional results on response fields* in *Supplemental Information*). Analysis of the error distribution demonstrates that increasing RF size decreases MSE by concentrating the distribution of squared-error closer to zero (Figure 3E), thus correcting the undesirable feature of randomly distributed errors in the full-order code case described above (see *Additional results on response fields in Supplemental Information*). In addition, increasing RF size decreases the required population size for mixed codes, and modestly increases the number of cases in which mixed codes outperform pure codes (Figure S3E, G). We also show increased noise robustness from mixed codes in simulations of a code for continuous stimuli under MSE, using continuous RFs (Figure S4A). Thus, mixed codes are an effective strategy for reliable and efficient coding not just for decision-making systems, but also in sensory systems – which is consistent with their widespread observation in sensory brain regions[30–35].

### Experimental evidence that mixed codes support reliable decoding

Previous theories about NMS have focused on the fact that it enables flexible linear decoding, and there is experimental evidence that the dimensionality expansion provided by NMS is linked to performance of complex cognitive behaviors[4, 27]. Here, we have shown that mixed codes also provide more general benefits for reliable and efficient information representation in the brain, independent of a particular task and without assuming linear decoding. Thus, our work predicts that mixed codes will be used widely in the brain, instead of being used only for features relevant to particular complex tasks.

To understand whether the brain exploits mixed codes for their general reliability and efficiency rather than only for their ability to enable flexible computation, we test whether the brain implements NMS when it would not enable the implementation of any behaviorally relevant linear decoders. To do so, we analyze data from a previously published experiment[45] that probed how two behaviorally and semantically independent features are encoded simultaneously by neurons in the lateral intraparietal area (LIP). In the experiment, monkeys performed a delayed match-to-category task in which they were required to categorize a sample visual motion stimulus (Figure 4A), and then remember the sample stimulus category to compare with the category of a test stimulus presented after a delay period (Figure 4B, top). In addition to the categorization and working memory demands of the task, the animals were also (on some trials) required to make a saccadic eye movement either toward or away from the neuron’s RF during the task’s delay period (Figure 4B, bottom, and see *Experimental details and task description* in *Methods*). Because LIP activity is known to encode information related to categorical decisions and saccades, this experiment characterized the relationship between the representation of these two features at the single neuron and population level. Despite the saccade being irrelevant to the monkey’s categorical decision in this task, LIP activity demonstrated both pure (*O* = 1, on average 43.5 % of the population tuned for each term, bootstrap test, *p* < .05) and mixed category and saccade (*O* = 2, on average 17.0 % tuned for each term, bootstrap test, *p* < .05) tuning (Figure 4D, E). This pattern of mixed and pure tuning is consistent with a composite code including RFs of multiple orders. Such codes have performance that falls between codes of either the lowest or highest included order alone, but their heterogene-ity may provide other benefits. Crucially, mixed codes also provide benefits when decoding only one of the two features at a time (Figure 4F). Thus, with two behaviorally and semantically independent features, the brain still implements a mixed code even though it does not enable the implementation of any behaviorally useful linear decoders. The mixed code does, however, improve the reliability and efficiency of the encoding, suggesting that the brain may explicitly utilize mixed codes for that purpose. Further, contemporaneous work has demonstrated that the bat is likely to exploit the reliability benefits of NMS for the coding of two-dimensional continuous head-direction information – as well as described reliability benefits of full-order mixed codes for continuous stimuli[31] (and see *Error-reduction by NMS in the continuous case in Supplemental Information*).

**Figure 4:**
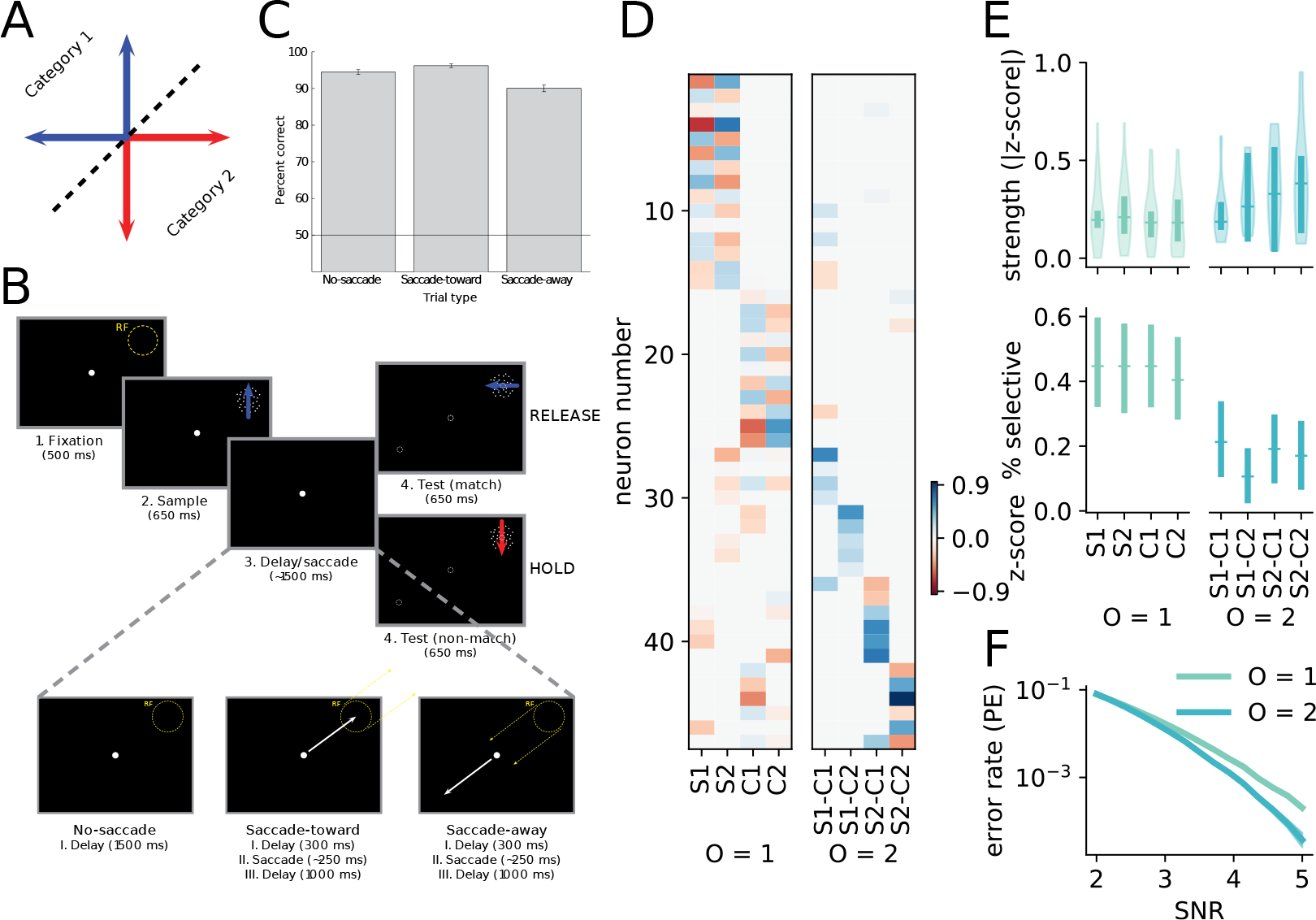
Mixed codes support reliable decoding in the brain, not only flexible computation. **A** The learned, arbitrary category boundary on motion direction used in the saccade DMC task. **B** A schematic of the saccade DMC task. **C** The average performance of the two animals on the saccade DMC task plotted for the different saccade conditions. **D** A heatmap of the z-scored magnitude of the coefficients for each term in the linear model. It is sorted by largest magnitude term, from left to right. The linear models were fit using the LASSO method and terms were tested for significance using a permutation test (*p* < .05), only neurons with at least one significant term were included in this and the following plots. **E** (top) The average strength of significant tuning for each term across the neural population, *O* = 1 tuning is on the left, and *O* = 2 tuning is on the right. (bottom) The proportion of neurons in the population that have pure selectivity (left) for the two saccade targets and two categories of motion and nonlinear mixed selectivity (right) for each of the four saccade target and category combinations. Error bars are bootstrapped 95 % confidence intervals. **F** Single-feature decoding performance for a code chosen to mirror the conditions of the task, with *K* = 2 and *n* = 2. Mixing features together is advantageous even when decoding those features separately.

## Discussion

We have shown that NMS and mixed codes are an effective and general strategy for reliable and efficient communication. Further, we have demonstrated that, rather than pure (*O* = 1) or fully-mixed (*O* = *K*) codes always providing the most reliable encoding, the optimal code order tends to lie between these two extremes (1 *< O < K*) depending on the number of stimulus features, the required fidelity of those features, and the number of neurons or total metabolic energy available to encode the information (Figure 2D). This set of intermediately mixed codes has not previously been analyzed in this context, despite likely being the dominant form of NMS that exists in the brain. Intermediately mixed codes may also have an important additional benefit. The representations produced by the full-order mixed code (*O* = *K*) may be difficult to learn and to generalize from[46], due to the fact that each response pattern is the same distance from all other response patterns. Intermediately mixed codes (1 ≤ *O* < *K*) ameliorate this by placing response patterns that are nearby in stimulus space nearby to each other in response space as well. That is, intermediately mixed codes carry more information about the stimulus space (rather than just the stimulus) in their responses than full-order codes, and this information may be crucial for behavioral performance and learning[47]. Lastly, we have shown experimental evidence that the brain implements mixed codes even when they do not facilitate behaviorally relevant linear decoding, but do improve the reliability and efficiency of encoding.

This work differs substantially from most previous work on optimal RFs in four principal ways. First, the dependence of code reliability on RF order, or dimensionality, has not been comprehensively described. We show that codes using RFs of intermediate dimension (1 *< O < K*) are most reliable in a wide variety of cases (Figure 2D), but these codes have not been previously studied outside of binary stimulus features. Second, we directly compute the probability and magnitude of errors for our codes rather than maximizing quantities with indirect relationships to error, such as Fisher information and mutual information. This reveals the performance of our codes even in low SNR regimes, where the indirect relationships are not guaranteed to provide correct descriptions of performance. Third, by accounting for the metabolic cost of both the total spike rate as well as the minimum population size required to implement each of our codes while keeping coverage of the stimulus space constant, we disentangled performance decreases due to a lack of coverage of the stimulus space from those due to the properties of the encoding itself. Fourth, we have investigated differences in code performance across different orders for both discrete and continuous stimuli as well as both binary (PE) and distance (MSE) error metrics. These different contexts have revealed several nuances, including that, for discrete stimuli, increasing RF size tends to increase PE, but decrease MSE – highlighting the ways in which RF shape and size can influence which kinds of coding errors are likely for different coding strategies, which has not received extensive study in neuroscience. Thus, this work provides a novel perspective on multiple understudied neural coding problems.

This work also ties directly to existing work in the experimental and theoretical neuroscience literature. Most centrally, we link the previously described flexible linear decoding benefits of NMS to considerations of reliability and efficiency in neural codes. Experimental work focusing on the utility of NMS for flexible linear decoding has already demonstrated the ubiquity of mixed codes in prefrontal cortex[4], as well as a putative link from NMS to behavior[27]. Theoretical work has demonstrated that random connectivity in recurrently connected neural network models produces a mixed code for stimulus features[48]. In addition, while prefrontal cortex has been shown to exhibit more NMS than expected purely due to those random connections, biologically plausible, unsupervised Hebbian-like plasticity rules applied to similar model networks increases the prevalence of NMS to levels consistent with those observed experimentally[49]. Thus, not only do mixed codes provide two substantial and separate benefits to the brain, they are also naturally produced by known neural phenomena – that is, they do not require fine tuning.

Our broader view of the benefits of NMS also helps to explain its observation in numerous systems aside from macaque prefrontal cortex. Recent experimental and theoretical work on the *Drosophila* larva demonstrates that NMS develops quickly in olfactory-input receiving Kenyon cells of the mushroom body. Further, the mixed code that is developed by the mushroom body produces fewer errors than random codes with similar orders, but that do not guarantee full coverage of the olfactory space[32]. Thus, mixed codes do not arise only as a product of random connections, but are likely refined to serve reliable communication early in development even in conditions with relatively small numbers (i.e., ~ 100) of neurons. Further, interrogation of the bat head-direction system has revealed a dynamic code, in which single neurons shift from having pure selectivity for long decoding intervals (e.g., during long-distance navigation) and mixed selectivity for short decoding intervals (e.g., during rapid maneuvering and chase behaviors). In a population of neurons with fixed size, this strategy is advantageous if adequate coverage of the space cannot be achieved by a mixed code[31]. Thus, mixed and pure codes must both be decodable by the brain, and it appears that the most reliable and efficient code is selected moment-to-moment as the time available for decoding shifts. More generally, mixed codes have been observed across diverse sensory and non-sensory systems[4, 27, 30–38, 50], indicating that their usefulness is not only due to enabling flexible linear decoding, but also due to their coding reliability and efficiency.

Our work also illustrates several tradeoffs that can be leveraged to understand the organization of feature representation across different brain regions. First, we have shown that the optimal code order decreases as the fidelity of the stimulus features (i.e., the number of values each feature takes on, *n*) increases. Thus, it will be important to directly compare the fidelity and code order across different levels of the sensory-processing cortical hierarchy. Second, our framework can give insight into parallel sensory processing pathways. In particular, large numbers of features (large *K*) quickly become impractical to represent with high order (*O*) mixed codes, due to an exponential increase in the required population size with order (Figure 1E, left). However, the brain may still be able to leverage some of the benefits of of mixed codes by representing the *K* features in two (or more) distinct subpopulations that each represent [*K/*2] features. The visual[51] and auditory[52] systems in macaques are both thought to be split into multiple streams representing distinct features, consistent with this idea. In particular, if mixed codes do arise naturally from unsupervised synaptic plasticity, then keeping the number of features represented in any particular brain region beneath some threshold may be an effective strategy for guaranteeing full coverage of the stimulus space.

Overall, our work has shown that NMS is an effective and practical strategy for reliable coding in the brain. Guaranteeing this reliability, in the face of unreliable neurons, is likely to have fundamentally shaped the functional and even anatomical architecture of neural systems. Developing an understanding of the role of code order, or RF dimensionality, in reliable and efficient coding will give insight into this much broader problem.

## Acknowledgements

We gratefully acknowledge Chris Rishel and Gang Huang for conducting the experiments which provided the neurophysiological data. We also thank Xaq Pitkow, Jeff Beck, Nicolas Masse, Jared Salisbury, Krithika Mohan, and Yang Zhou for their comments on and useful discussion of earlier versions of this manuscript. This work was supported by NIH F31EY029155 (WJJ), NSF CAREER-1652617 (SEP), NIH R01EY019041(DJF), CRCNS NIH R01MH115555 (DJF), NSF NCS 1631571 (DJF), and a DOD Vannevar Bush Fellowship (DJF).

## Author contributions

WJJ conceived of the project. SEP and DJF supervised the project development. WJJ created the model and performed the calculations, model simulations, and data analysis. DJF designed and supervised the experimental work. SEP supervised the theoretical work. WJJ, SEP, and DJF wrote the paper.

## 1 Methods

### 1.1 Definition of the stimuli

In defining our stimuli, we make two assumptions. First, we assume that our stimuli are described by *K* independent features. This is equivalent to assuming that our stimuli were pre-processed by an efficient coding procedure that isolated the independent components[1] of our stimulus set[2]. Second, we assume that the stimulus features are discrete and uniformly distributed; this simplifies our mathematical analysis, but also makes our analysis relevant to cognitive, categorical representations. In addition, simulations with continuous stimulus features have qualitatively replicated our core results (see *Error-reduction by NMS in the continuous case in Supplemental Information*).

Thus, a stimulus is represented by a vector of *K* discrete values. Each value corresponds to one of the *K* independent features of the stimuli. The nature of the value object does not matter, we only require that it is possible to decide whether two values corresponding to the same feature are equal. For a stimulus *x* with *K* features,

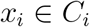

for *i* ∈ [1, …, *K*], where *C*_*i*_ is the set (of size *n*_*i*_) of all possible values for feature *i*. Using the equality function, we implement an indicator function,

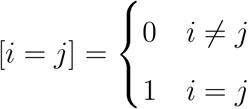

for all values of all features. From our assumption that the stimuli are composed of independent features that take on discrete values with a uniform probability, it follows that all value combinations are valid, giving *M* = ∏_*i*_*n*_*i*_ possible stimuli, and equally probably, so 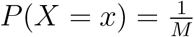.

### 1.2 Definition of the codes

Our definition for nonlinear mixed selectivity (NMS) follows that given in [3]. We describe it with some generalizations here.

The codeword *c* corresponding to stimulus *x ∈ X* is produced by

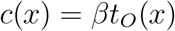

where *β* is a matrix of size *N × D* and *t*_*O*_ (*x*) is the encoding function of order *O*. Our codes will primarily be differentiated by *t*_*O*_ (*x*), while *β* will be used to equalize their representation energy *V* and population size *N*.

The elements of the vector *t*_*O*_ (*x*) are products of indicator functions, and therefore can only be either one or zero. In particular, for order *O*, the vector *t*_*O*_ (*x*) contains one element for the product of each valid combinations of feature-value indicator functions of size *O*. More formally, for 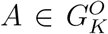where 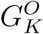 is the set of all combinations of *O* elements from [1, …, *K*], and 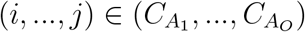,

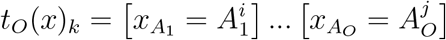

with individual neurons (indexed by *k*) corresponding to all feature combinations *A* and all value combinations for those features (*i*, …, *j*).

Thus, 1 ≤ *O* ≤ *K*, where *K* is the total number of stimulus features, and all codes with *O* ≥ 2 are mixed while codes with *O* = 1 are pure codes, following [3]. We will use the term “neuron” to refer to coding units in our models and simulations as well as to refer to biological neurons in the brain to make their analogous roles clear. In our formulation, both mixed and pure codes will always have complete coverage; that is, there will be a neuron coding for every feature value or possible combination of feature values and each of the *M* stimuli will have a corresponding unique codeword.

#### 1.2.1 Code example

For *K* = 3 and *n* = 2, under our formalization there are codes of three different orders that code for the *n*^*K*^ stimuli.

**O = 1**: This code has *nK* neurons and below we give some example stimuli (on the left, with the three features each taking on one of their two possible values, 1 or 2) and codewords (across the activity of the neurons, on the right):

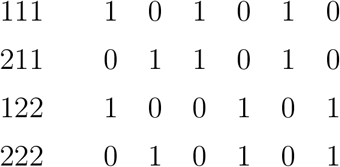

Note that for each of these stimuli, there are always three neurons responding with 1. Further, the smallest distance between any two codewords is 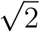, between 111 and 211 as well as 122 and 222. This is of course not the smallest number of neurons that we could use to represent the set of 8 stimuli. The smallest number of neurons that could represent these stimuli is log_2_ *n*^*K*^ = log_2_ 8 = 3 neurons, which could use a representation similar to the one we have used to represent the stimuli on the lefthand side of the above table.

Thus, this encoding strategy has added redundancy to our representation of the stimuli.

**O = 2** This code has 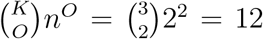 neurons. It can be viewed as three separate *O* = 2 codes for the three different size 2 subsets of the 3 features. We make that explicit in our example:

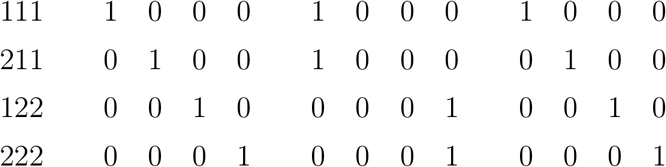

Note that any two of these three subpopulations alone would produce a code with unique codewords for each of the stimuli. However, they would preferentially represent one of the three features and cause errors to be more likely for the other two features. The minimum distance between any of the stimuli is now 2 and the number of neurons active is 3.

**O = 3**: This code has *n*^*K*^ = 8 neurons, that each code for a unique combination of the three features – and therefore for a unique stimulus. As in:

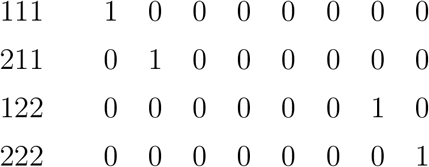

Note that there is now only one neuron active for each stimulus, and the minimum distance is 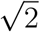.

Next, we formalize these properties: population size, minimum distance, and representation energy (or the number of active neurons) and derive expressions for each of them for general *K* and *n*.

### 1.3 Code properties

#### 1.3.1 Population size (*D*_*O*_) of the codes

The population size of a code is the length of *t*_*O*_ (*x*) for that code. Since we know that a code of order *O* will have an element for each possible combination of feature-values of size *O*, the length of the vector can be framed as a counting problem:

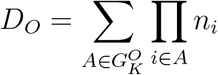

where 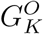 is the set of all subsets of [1, …, *K*] with size *O* and *n*_*i*_ = |*C*_*i*_|. This expression is somewhat cumbersome, so, for ease (and without affecting our results), we assume that *n* = *n*_*j*_ for all *j ∈* [1, …, *K*]. This gives,

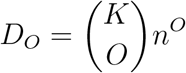

where 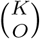 is the binomial coefficient, defined as

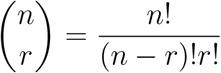

if *n ≥ r*, otherwise 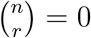.

For *O* = 1 (the pure code), the population size is

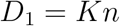

and, for *O* = *K* (the fully mixed code), it is

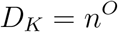

Thus, the population size (i.e., the length of the vector) grows exponentially with the order of the code.

#### 1.3.2 Representation energy (*P*_*O*_) of the codes

We quantify the amount of energy that each coding scheme uses to transmit codewords. In particular, we will model energy in two ways and will see that these are equivalent for large *n*_*i*_ and do not substantially change our results for smaller *n*_*i*_.

##### Sum of variance across dimensions

Taking the variance of a particular dimension as the energy used by that dimension for coding, we can simply take the sum across all of the dimensions. With the definition of variance,

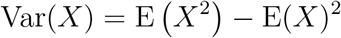

we can express representation energy (*P_O_*) as:

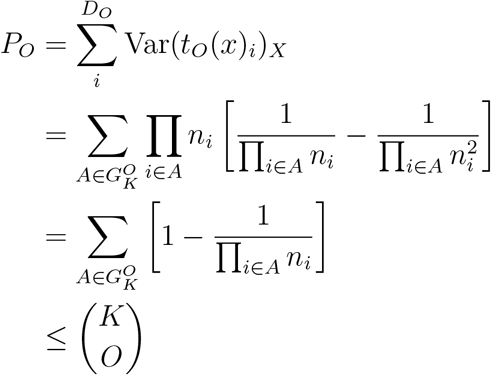

With large *n*_*i*_ the second term in the sum becomes very small, and we can see that the upper bound of the last line gives a good approximation of the representation energy.

So, for *O* = 1,

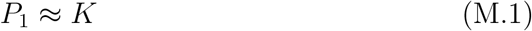

For, *O* = *K*,

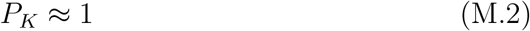

##### Points on a sphere

We also notice that for a code of a particular order, all of the codewords lie on a high-dimensional sphere with radius *w*_*O*_. The radius of this sphere provides a different notion of energy consumption, through 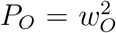. Formally, it differs from the notion of energy consumption given above only in that the squared mean activity is not subtracted. That is,

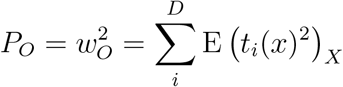

rather than

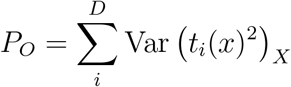

Following the derivation above, the radius is:

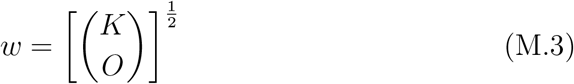

and the representation energy is

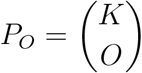

and, for *O* = 1,

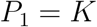

and, for *O* = *K*,

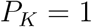

That is, this gives the same answer as our other measure for energy, but does not depend on the assumption that the *n*_*i*_ are large.

Use of either of these two measures does not substantively affect our results. In our simulations, we will use the former because it slightly benefits pure codes (because the mean activity of neurons in pure codes is generally higher than that in mixed codes, so there is a larger reduction in their representation energy by the subtraction of the squared mean) and we are exploring the benefits of mixed codes.

#### 1.3.3 Minimum distance (Δ) of the codes

The smallest distance between any two codewords is directly related to the probability that a decoder will make an error when attempting to discriminate between those two codewords, and can be used to bound the performance of decoders in general.

##### Statement 1.

*The distance between two stimulus codewords is given by*

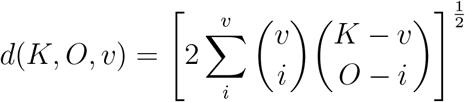

*where v is the number of features the stimuli differ in, O is the order of the code, and K is the number of features*.

*Derivation.* Using the set 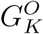 with 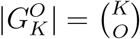, we see that when we change a feature *i* ∈ [1, …, *K*], by the definition of the indicator function and of our codes, we know that one term (a product of indicator functions) in each feature combination that includes *i* will flip from 0 to 1 and another term will flip from 1 to 0. Thus, given the subset 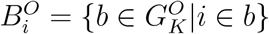, we obtain a distance of 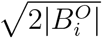 from changing the value of feature *i*. When we change a second term, *j*, we obtain 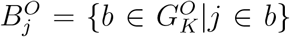 The distance between the two stimuli is then related to the size of the union of these two sets: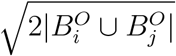.

So, to find the distance between two codewords, we need to count the number of features in which they differ and then find the distance, given the order of the code *O* and the number of stimulus features *K*.

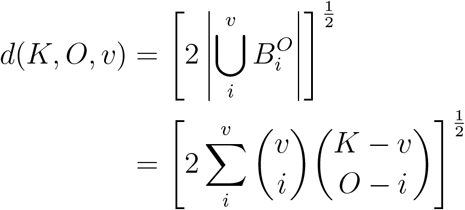

where the second binomial coefficient counts the number of subsets containing exactly *i* of the *v* changed features and the first binomial coefficient counts the number of different ways *i* features could be chosen from the *v* changed features. Since our codes include all combinations, the identities of the features changed does not matter – only the number of them.

Next, it will be useful to know that this distance function is increasing with *v*, as, combined with statement 1, it will allow us to find the minimum distance.

##### Statement 2.

*The function d*(*K, O, v*) *is increasing with v*.

*Derivation.* We want to show that *d*(*K*, *O*, *v*) ≤ *d*(*K*, *O*, *v* + 1).

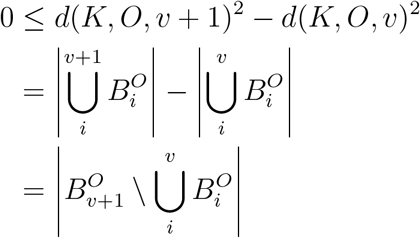

where the last line is the size of the set of values that are in 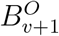 and not in any of the other 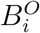 for *i* ∈ [1, …, *v*]. The relationship holds because a set cannot have a negative size. Thus, *d*(*K*, *O*, *v* + 1) ≥ *d*(*K*, *O*, *v*) and therefore the function *d* is increasing in *v*.

Now, we can find the minimum distance using both of the statements above.

##### Statement 3.

*The minimum distance of a code of order O for stimuli with K features is given by*

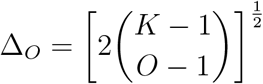

*Derivation.* By statement 2, we know that the minimum value of *d*(*K*, *O*, *v*) occurs when *v* = 1. We can then evaluate our expression for distance, found in statement 1, at *v* = 1

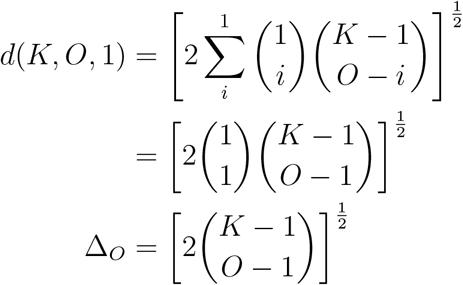

Now we can evaluate this expression for any *K* and *O* that we desire. For *O* = 1,

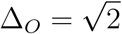

and, for *O* = *K*,

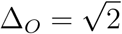

While the minimum distance for these two codes is the same, their representation energy is different (see Eq. M.1 and Eq. M.2). Further, minimum distance and power are both weakly unimodal around *O* = *K/*2 (Figure 1e, center and right).

### 1.4 Minimum distance-representation energy ratio

A straightforward way to describe code performance in a single number is to take the ratio between minimum distance and representation energy. Codes with larger ratios will typically have a lower probability of decoding error given the same noise level.

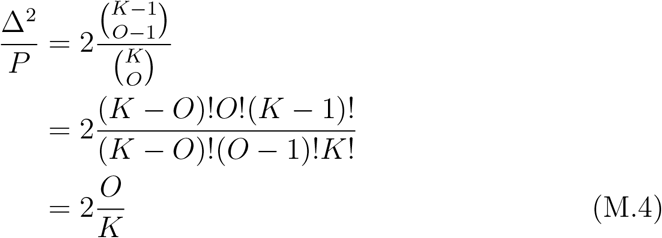

which is strictly increasing with order (Figure 1f, left).

### 1.5 The amplifying linear transform (*β*)

To compare between codes, we use *β*, a matrix, to perform a linear transform of the codewords *t*_*O*_ (*x*) that can be chosen to increase or decrease representation energy *P*_*O*_ to a fixed value *V* as well as increase (but not decrease) population size *D*_*O*_ to a fixed value *N*. Thus, we can compare codes of different orders that have the same representation energy (*V*) and population size (*N*).

In choosing *β*, we must satisfy four constraints:

1. *N* ≥ *D*
2. *β*^†^*β* = *I*, where *I* is the *D* × *D* identity matrix and *β*^†^ is the pseudoinverse of *β*.
3. The vector length, *H*, of each column in *β* must be the same.
4. E(*β*_*ij*_ *β*_*ik*_)_*j≠k*_ = 0; this will be true, for instance, for *β* where the rows or columns are sampled from a Normal distribution with a covariance matrix that is proportional to the identity matrix.

This flexibility in the choice of *β* can also be used to produce more granded rather than strictly binary responses to stimuli across our neural populations. However, as we will show, it is only the vector length of the columns of *β*, *H*, that affects the performance of the code.

#### Statement 4.

*For β with length H, the amplified code βt_O_* (*x*)*, where t_O_* (*x*) *has representation energy P_O_, will have representation energy H*^2^*P_O_.*

*Derivation.* We prove this using the points on a sphere definition of energy. So, the energy of the original code *t*_*O*_ (*x*) is given by,

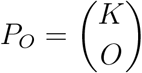

After applying *β*, we want to find the square of the average distance of the codewords from the origin, or *V*, under the points on a sphere definition of energy.

So, we want to find, where *c*(*x*) = *βt_O_* (*x*) and *X* is the set of stimuli,

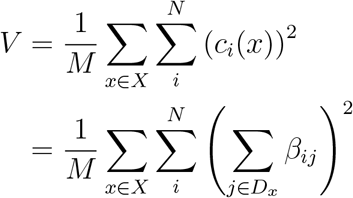

where *D_x_* is the set of non-zero indices of *t*_*O*_ (*x*) for *x*

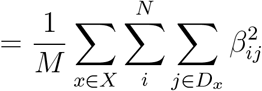

by the definition of *β*, constraint 4

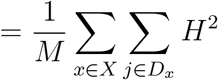

by the definiti n of *β*, constraint 3

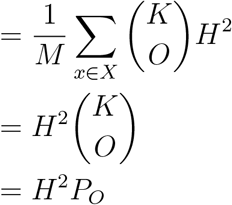

Thus, we can give different codes the same representation energy *V* by choosing 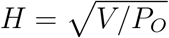 for each *O*.

#### 1.5.1 The effect of *β* on minimum distance

##### Statement 5.

*The distance between two points c_i_* = *βt_O_* (*x_i_*) *and c_j_* =*βt_O_* (*x_j_*), *represented as* 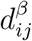, *is given by*

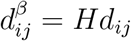

*where d_ij_ is the distance between the two points t_O_* (*x_i_*) *and t_O_* (*x_j_*).

*Derivation.* We know that points *x_i_* and *x_j_* are both 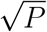 units away from the origin while codewords *c_i_* and *c_j_* are 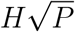 units from the origin (by statement 4 and equation M.3). We want to find 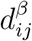.

The angle between the two points is

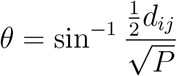

so, we can rearrange to find:

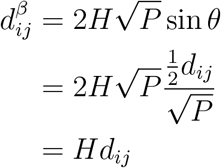

It follows directly from statement 5 that the minimum distance after *β* is applied, *δ*, is given by

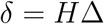

Further, it follows that the ratio given in Eq. M.4 is not altered by *H*, or choice of particular *β*, since

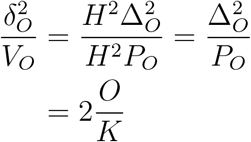

### 1.6 Full channel details

We simulated codes of all possible orders for particular choices of *K* and *n*. Three important choices were made for these simulations. First, the codewords from each code were passed through a linear transform *β*. The linear transform was used to equate the population size and representation energy of different order codes, such that we could investigate code performance when each order of code had the same number of participating units and the same signal-to-noise ratio (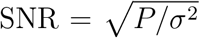 where *σ*^2^ is the noise variance), as in Figure 2 and see *The amplifying linear transform (β)* in *Supplemental Information*. Second, the noise in the channel was chosen to be additive and to follow an independent Normal distribution across code dimensions. Third, we use maximum likelihood decoding (MLD) to estimate the original stimulus. This choice is consistent with Bayesian and probabilistic formulations of neural encoding and decoding[4–6]. While inclusion of noise correlations would be an interesting topic for future research, we show here that they are not essential for any performance increases due to NMS.

#### 1.6.1 Code availability

All of the code for the simulations was written in Python (3.6.4) using NumPy (1.14.2), SciPy (1.0.1)[7], and Scikit-learn (0.18.1)[8]. The code is available on request. For each SNR and each code order, approximately 5000 trials were simulated.

### 1.7 Union bound estimate

While the minimum distance-representation energy ratio we derive in Eq. M.4 provides useful insight into the performance of codes of different orders, it does not give a direct estimate of the probability of decoding error. In particular, it is difficult to interpret the magnitude of performance differences without incorporating the magnitude of the noise itself, the decoder used, and the arrangement of all of the codewords in coding space to estimate PE directly. Here, we incorporate the details of the full channel to directly estimate PE via a union bound estimate (UBE).

That is, with the channel,

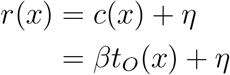

where *η*~N(0*, σ*^2^) (see Figure 1a for a schematic) and a maximum likelihood decoding function *f* such that 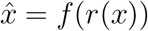 where 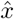 is the maximum likelihood estimate of *x* given *r*(*x*), we want to estimate the probability that 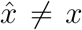 across *X*. To begin,

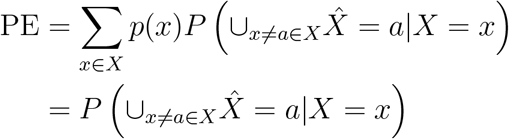

by statement 7

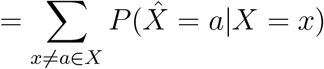

by the disjoint nature of decoding events

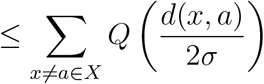

where *Q*(*y*) is the ccdf at *y* of 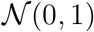 and *d*(*x, y*) is the Euclidean distance between the *codewords* corresponding to *x* and *y* (i.e., the Euclidean distance between *βt_O_* (*x*) and *βt_O_* (*y*)).

From here, there are several different ways to approximate (or upper bound) PE. We will focus on a nearest neighbor approximation, where we assume that the majority of the PE arises from errors made to the incorrect codewords nearest to the correct codeword (i.e., the nearest neighbors). That is, stimuli *a* with *d*(*x, a*) = *δ_O_*. This works as an approximation due to the exponential decrease of *Q*(*y*) with *y* (and see Figure 2 for empirical confirmation).

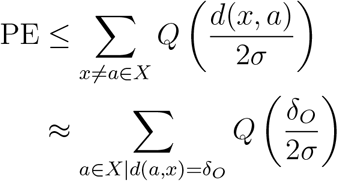

taking only the nearest neighbors

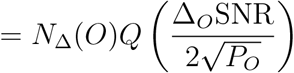

applying Eq. M.4

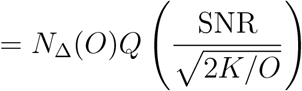

where *N*_Δ_(*O*) is the number of nearest neighbors the code of order *O* has, we derive it in *Code neighbors* in *Supplemental Information*. Thus, we can see that PE depends most strongly on the minimum distance-representation energy ratio and SNR, but also depends on the number of nearest neighbors a particular code has at minimum distance. This number is the same (Eq. S.1) for codes with 1 ≤ *O* < *K*, but vastly increases for codes with *O* = *K*.

### 1.8 Total energy

Similar to [9], we assume that all neurons, whether spiking or not, consume some baseline, non-zero amount of energy – due to passive maintenance processes, including the circulation of ion channels, and due to spontaneous activity. We define this amount of energy to be equal to one unit. Next, we assume that spiking neurons consume the baseline energy plus an amount of energy proportional to the square of their firing activity; this activity summed across the population is the representation energy (*P_O_*). So, the total energy consumption of a code, *E*, can be written:

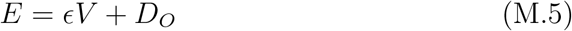

where *ϵ* controls the proportional cost of spiking relative to passive maintenance costs. This *ϵ* will vary between neuron types, but has been estimated by experiment to be around 10 to 10^2^[9].

From Eq. M.5, we see that a code of order *O* allocated *E* total energy would have,

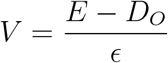

and

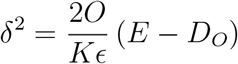

where only codes with *V >* 0 (that is, *E > D_O_*) can be implemented in practice. This *δ* is used in the comparisons for Figure 2d. From this expression, we observe that the particular value of *ϵ* does not change the relative performance of codes with different orders. So, our results in Figure 2d do not depend on *ϵ*.

Further, we find that when *δ_O_* = *δ*_*O*+1_ as a function of *E* to discover when the *O* + 1-order code will begin to outperform the *O*-order code:

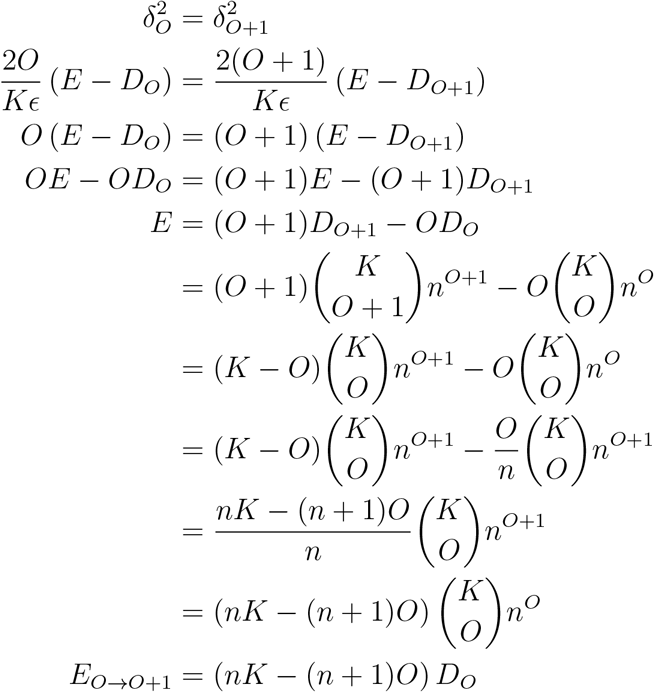

and using this for *O* = 1, we find that

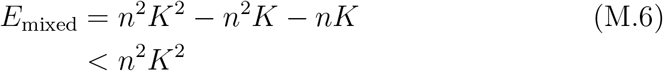

such that for *E > E*_mixed_ a mixed code (i.e., a code of order *O >* 1) will always provide better performance than a pure code.

### 1.9 Experimental details and task description

We used experimental data in Figure 4 that was previously published in a separate study [10]. The full methods are given in the original paper, though we briefly review several key points here. The data may be requested from the authors of the previous study.

#### 1.9.1 The behavioral task

See the schematic in Figure 4b. First, a moving dot stimulus in a direction that was on one side of a learned category boundary was presented while the animal fixated. Then, there was a delay period during which the animal was compelled to saccade to one of two locations before, finally, a second motion stimulus was presented and the animal reported whether the category of the first (or sample) stimulus matched the category of the second (or test) stimulus. The division of the 360° of motion direction into two contiguous categories was arbitrary, and learned by the animals over extensive training.

#### 1.9.2 The electrophysiological recordings and analysis

The experimenters recorded from 64 lateral intraparietal area (LIP) neurons in two monkeys (monkey J: *n* = 35; monkey M: *n* = 29) during performance of the DMC task. Recordings were performed using single 75 µm tungsten microelectrodes (FHC). Units were sorted offline, and selected for quality and stability. No information about the LIP subdivision from which each neuron was collected is available.

Linear models for motion category (category 1 or 2) and saccade direction (toward or away from the neuronal RF) with interaction terms (between category and saccade direction) were fit using an L1 prior in scikit-learn (i.e., the Lasso fitting procedure) to all neurons with greater than 15 trials for each of the four conditions. Coefficients were tested for significance via a permutation test at the *p* < .05 level. Spikes were counted in the 20 ms to 170 ms window after the saccade was made and then spike counts for each neuron were z-scored across the four conditions.

### 1.10 The rate-distortion bound and mutual information calculation

To calculate the rate-distortion bound (RDB) for our source distribution, we use a Python implementation of the iterative Blahut-Arimoto algorithm[11, 12]. Since the optimization problem is convex, the algorithm is guaranteed to converge on the right solution, given enough iterations. To ensure an adequate number of iterations, we terminate the algorithm only when successive steps are less than 10^*−*10^ change in error probability magnitude.

To evaluate our codes alongside the RDB, we must calculate the mutual information between the stimulus distribution *X* and the distribution of our stimulus estimates 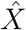 So,

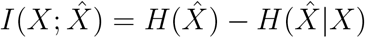

where

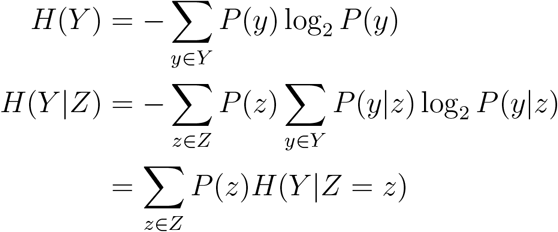

To compute these quantities, we rely the observation that 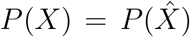 That is, both distributions are uniform, with 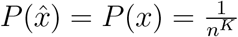. This can be seen from the fact that none of our codewords have more (or fewer) neighbors at any given distance than any of our other codewords (see statement 7).

Using this,

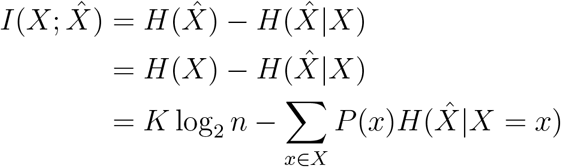

Since 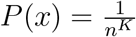 and 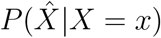 has the same entropy for all *x*, following from the observation above, it is enough to estimate

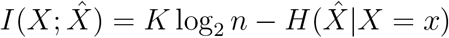

for a particular *x*. We do this via numerical simulations (see *Full channel details in Methods* for details).

## A Supplemental Information

### A.1 Glossary of terms

M: The number of stimuli transmitted by a code.
Δ_*O*_: The minimum distance of the code of order *O*.
*δ_O_*: The minimum distance of a code of order *O* after *β* is applied.
*P_O_*: The representation energy used by a code of order *O*.
*V*: The representation energy used by a code after *β* is applied.
*D_O_*: The population size of a code of order *O*.
*N*: The population size of the code after *β* is applied.
*K*: The number of features that a stimulus has.
*C_i_*: The set of values that feature *i* can take on.
*n*_*i*_: The size of set *C_i_*; that is, *n_i_* = *|C_i_|*
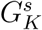: The set of all possible subsets of [1*, …, K*] with size *s*; {*X* ⊂ [1, …, *K*] : |*X*| = *s*}
*x*: A stimulus; a vector of length *K*, where *x_i_ ∈ C_i_* for all *i*.
*t*_*O*_ (*x*): The encoding function of order *O*. It takes a stimulus (*x*) and produces the representation of that stimulus in a code of order *O* – also referred to as the codeword. The representation is a vector of length *D_O_* of ones and zeros.
*β*: The amplifying transform. It is applied to the codeword (*t*_*O*_ (*x*)) and produces the amplified encoding; *β* is a matrix of size *N × D* and must satisfy the constraints given in *The amplifying linear transform (β) in Methods*.
*H*: The power in each column of *β*; 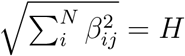 for all *j*.
*η*: A noise term. Here, always Gaussian, with *η ∼* N(0*, σ*^2^).
*c*(*x*): The amplified codeword corresponding to a given stimulus, *c*(*x*) = *βt_O_* (*x*). It is a vector of length *N*.
*r*(*x*): The noisy amplified codeword corresponding to a given stimulus, *r*(*x*) = *c*(*x*) + *η*. It is a vector of length *N*.
*f*(*r*): The maximum likelihood decoding function for a particular code. It solves the equation argmax_*x*_ *P* (*r|x*)*P* (*x*)*/P* (*r*).
*x*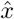: The estimate of *x*, derived from a noisy representation, 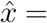 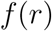.

### A.2 Code neighbors

For the UBE, it becomes necessary to know the number of codewords at minimum distance from any given codeword (*N*_Δ_(*O*)).

#### Statement 6.

*The number of neighbors at minimum distance for a code of order O N*_Δ_(*O*) *is given by:*

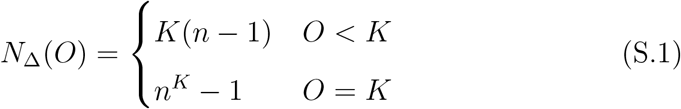

*Derivation.* From the fact that the distance function is increasing with *v* (statement 2), we know that *d*(*K*, *O*, 1) is the minimum of *d*(*K*, *O*, *v*), but it may or may not be a unique minimum.

Thus, we want to find *O* such that *d*(*K*, *O*, 1) *< d*(*K*, *O*, 2),

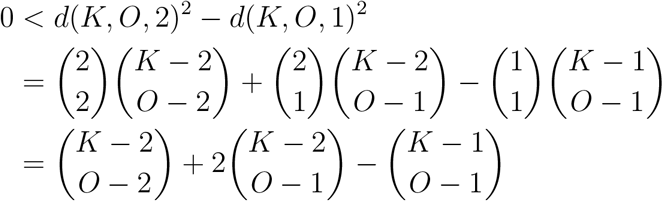

exploiting binomial identities to make all binomial terms equal

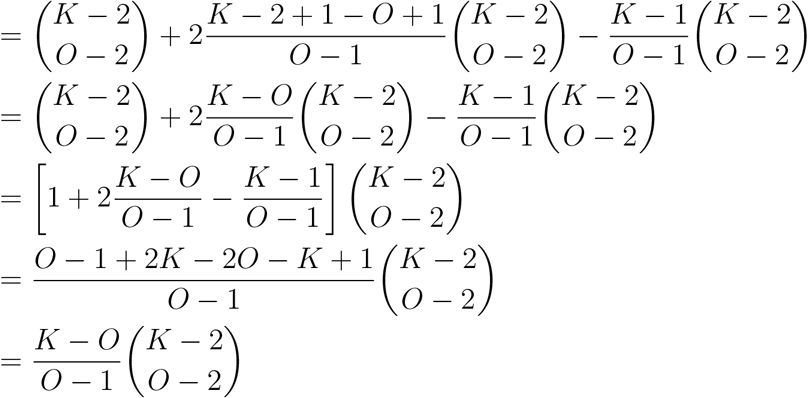

this is undefined for *O* = 1, which is undesirable

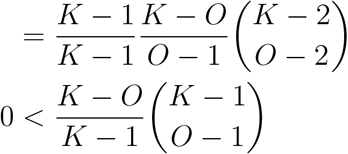

This last expression is true when 1 ≤ *O* < *K* and false otherwise (i.e., when *O* = *K*). When it is true, it implies that changing one stimulus feature produces codewords at a closer distance than changing two stimulus features. Now, we must find how many stimuli differ by a single feature from a given stimulus. Any single feature of the *K* features could be changed, and it could be changed to any one of *n −* 1 different values (excluding its current value) – so, *N*_Δ_(*O*) = *K*(*n −* 1) for *O* < *K*.

If *O* = *K*, then 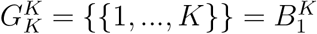 and since 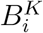 cannot grow beyond the size of 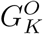 all codewords must be at the same distance. Thus, *N*_Δ_(*O*) = *n*^*K*^ − 1 for *O* = *K*.

#### Statement 7.

*The number of neighbors at a fixed distance does not depend on codeword identity*.

*Derivation.* We assume that the number of neighbors at a fixed distance does depend on codeword identity and show that this leads to a contradiction. We know that codeword distance does not depend on original codeword identity (statement 1), but does depend on the number of features that the stimuli differ by. Thus, for a set of codewords to have more neighbors at a particular distance than a different set of codewords, the corresponding set of stimuli must be able to differ in more ways from the corresponding set of other stimuli. Stimuli can differ by changing 1 to *K* of their *K* features to one of the *n* − 1 different values for each feature *C_i_*. For a set of stimuli to be able to differ in more ways than a different set of stimuli, that set of stimuli must have either more features or more possible values for each feature. Either of these would contradict our definition of the stimuli (see *Definition of the stimuli in Methods*).

### A.3 L1 norm for representation energy

To this point, we have used the L2-norm to characterize the relationship of spiking activity across the population to metabolic energy consumption in the form of representation energy. This is following decades of literature on neural coding[1] and communication theory[2]. However, there is some evidence to suggest that the L1-norm may be more appropriate for use in the brain[3]. Given the L1-norm, the distance to representation energy ratio from *Minimum distance-representation energy ratio* in *Methods* becomes:

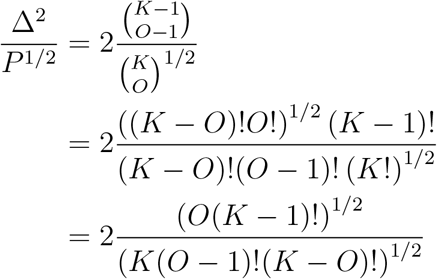

which is difficult to interpret intuitively, although one can observe that the *O* = *K* code always has a higher ratio than the *O* = 1 code, indicating that the broad intuition we have gained from our previous analysis holds here as well. However, codes that fall between these two extremes often provide superior performance to the *O* = *K* code. We demonstrate this in Figure S1.

### A.4 Poisson noise

To this point, the noise in our neural channel has been Gaussian distributed, which allows us to vary the SNR down our channel independently of representation energy or firing rate. However, neural firing rates are often viewed, at least roughly, as following a Poisson process, which implies a particular SNR at different firing rates due to a strict relationship between mean firing rate and firing rate variance (though experimentally observed firing rate-SNR relationships have not followed the one expected from a Poisson process[4]). Thus, it is possible that due to the different firing rates of individual neurons used in our codes (as only the sum firing rate is held constant across codes), Poisson noise could change which code performs best. However, in Figure S2, we show via simulations of our channel with Poisson instead of Gaussian variability that mixed codes still outperform pure codes.

### A.5 Additional results on response fields

Generalizing our current framework to allow flexibly sized response fields (RFs) requires only a reformulation of the indicator function. Instead of performing an equality operation, it should instead perform a set membership operation, as

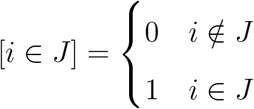

where the set *J* is, in this case, a contiguous sequence of feature values of length *σ*_rf_. Following this, for our main results, *σ*_rf_ = 1. Now, we explore how choosing *σ*_rf_ > 1 changes our results.

#### A.5.1 Effects on minimum distance, representation energy, and population size

Population size and representation energy change with RF size to ensure that full coverage of the stimulus set is maintained. To achieve this, we arrange the code dimensions in a series of *σ*_rf_ overlapping lattices, where each lattice has non-overlapping RFs in a grid pattern. This strategy is not guaranteed to be the most efficient tiling of the space, but it is simple to implement and analyze – and it approximately meets the theoretical estimate of the dimensionality of the most efficient tiling[5].

##### Dimensionality

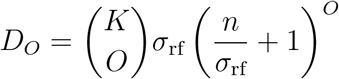

##### Power

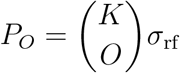

##### Minimum distance

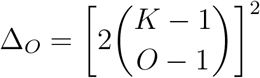

Note that minimum distance is not affected.

#### A.5.2 The optimal *σ*_rf_ for a given total energy

For a fixed *K*, *O*, *n*, and *E*, we want to find the *σ*_rf_ that maximizes minimum distance. For *E* = *ϵV* + *D_O_*, and using *δ*(*K, O, σ*_rf_*, V*) as an expression for minimum distance after application of *β* to produce a code with power *V*, we can write the problem as:

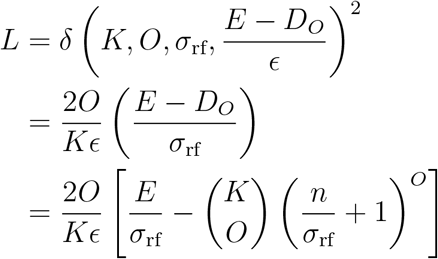

and now, to find the maximum, we will take the derivative 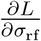,

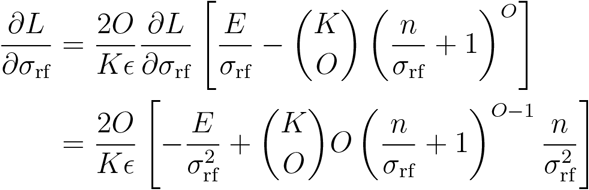

and now setting the LHS to zero,

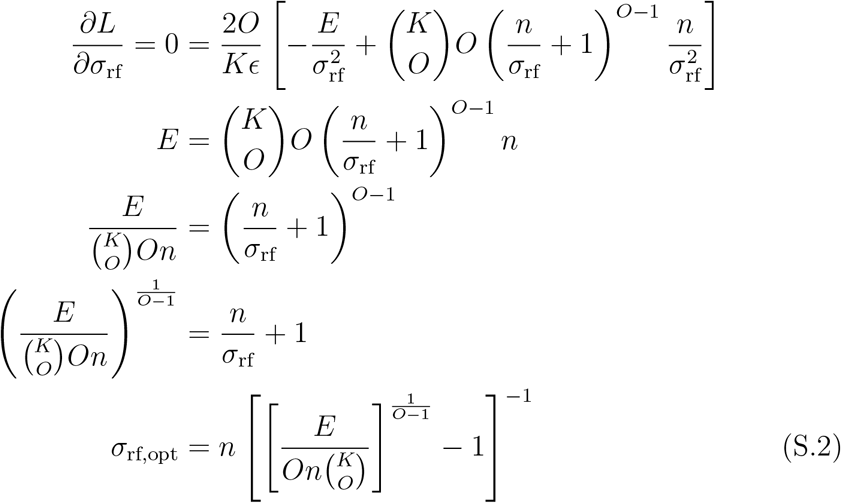

See Figure S3F for a plot of this function. This formalization does ignore benefits of *σ*_rf,opt_ > 1 for reducing the number of nearest neighbors of high order codes.

#### A.5.3 Effects on error distribution

Increasing RF size has the effect of pulling the distribution of squared-error distortion more concentrated toward zero, while increasing the overall probability of an error (see Figure 3D). The increase in overall probability of an error for fixed SNR can be understood by the expression for code power given above, where an increase in RF size increases the power consumption of the code without producing a change in minimum distance.

However, increasing RF size does produce a change in the number of codewords at minimum distance and at succeeding distances. To see this, we can focus on the *O* = *K* case: with *σ*_rf_ = 1, we know that all other codewords are nearest neighbors to a given codeword (Eq. S.1) because only one dimension is active for each codeword. If, instead, we have *σ*_rf_ = 2, we know that each RF has a volume of 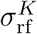 feature values, but their intersection must be of size 1. Thus, either active RF can be changed to 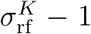 different RFs to still form a valid codeword. Thus, the number of nearest neighbors is 2(2^*K*^ − 1). With *σ*_rf_ = 2, all stimuli except the nearest neighbors will be at the same, further distance.

### A.6 Error-reduction by NMS in the continuous case

Here, we adopt continuous stimulus features and RFs to test how well the benefits of mixed codes generalize to the continuous case (also see [6] for a deeper investigation of the continuous case). In particular, with stimuli *x ∈ X* composed of *K* features, *x_i_ ~ U* (0*, n_i_*). Instead of the flat, discrete RFs defined in *Additional results on response fields* in *Supplemental Information*, we use Gaussian RFs,

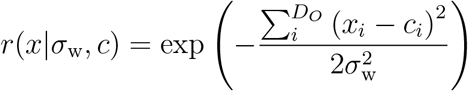

which are then scaled by the amplifying transform *β* as described in *The amplifying linear transform (β) in Methods*. The rest of the channel is identical to the channel described previously, including the additive noise. RFs are tiled in the same way, though now their width *σ*_w_ is independent of *σ*_rf_, which dictates their tiling – as in *Additional results on response fields* in *Supplemental Information*.

Our simulations show similar results to the discrete case (Figure S4), with higher order codes yielding lower MSE across all of the SNRs we investigated. Thus, the broad advantage of mixed codes apply in the continuous case as well. However, increasing RF size produces higher MSE, which is the opposite of our results in the discrete case. Future work is needed to discover why this is, and in what other ways the continuous case differs from the discrete case.

---

**Figure S1:**
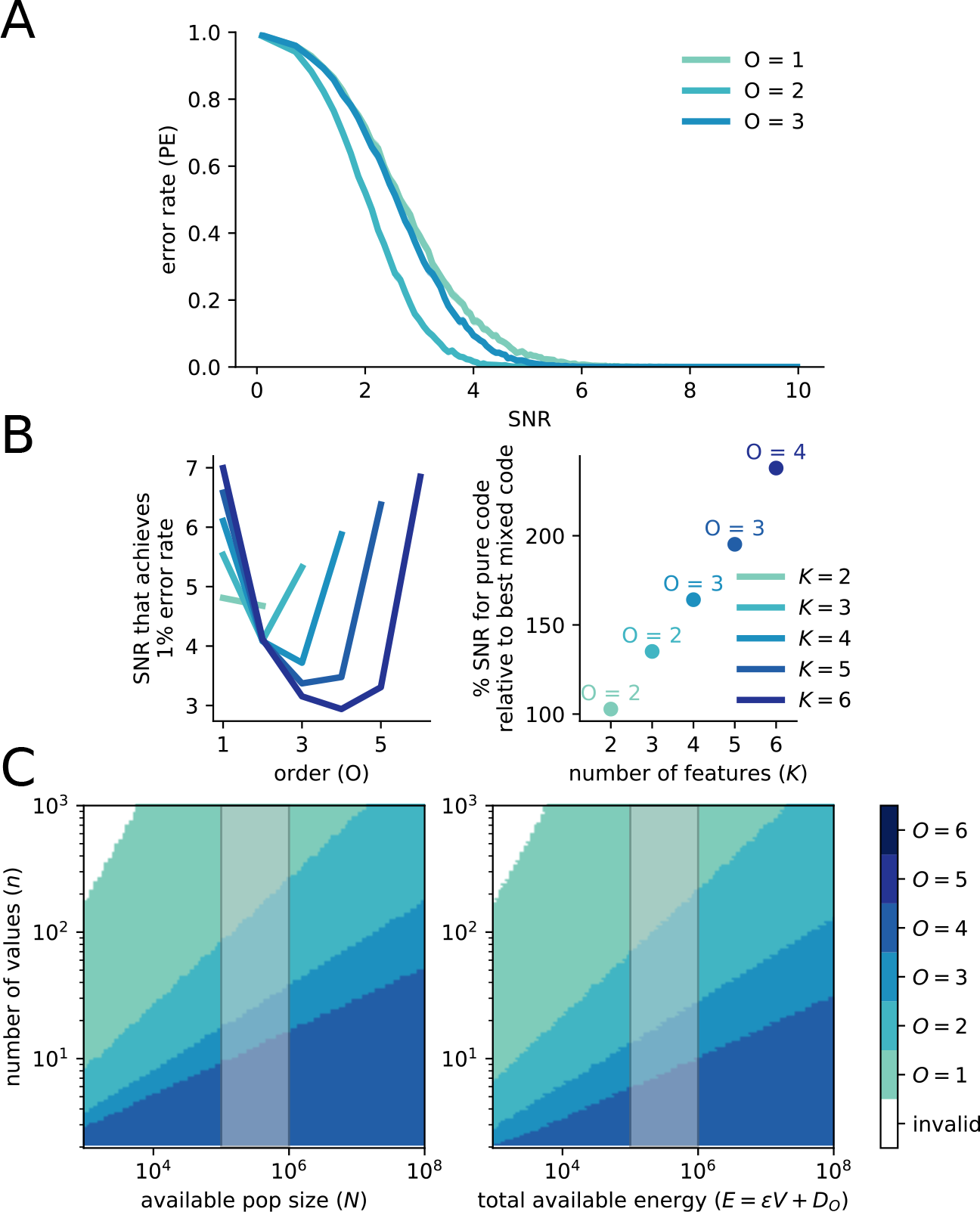
Using an L1 norm instead of an L2 norm to account for representation energy increases the performance of 1 *< O < K* codes, related to Figure 2 using the L1 instead of L2 norm for representation energy. **A** Simulation of codes with *O* = 1, 2, 3 for *K* = 3 and *n* = 5. **B** (left) Using the UBE, we show that for different *K* (with *n* = 5) the SNR required to reach 1 % decoding error tends to have its minimum around *K/*2. (right) The representation energy required by the pure code relative to that required by the best mixed code (given by point color and label) to reach 1 % decoding error. **C** (left) Given a pool of neurons with fixed size, the color corresponding to the code producing the highest minimum distance is shown in the heat map. The shaded area delineates the order of magnitude of the number of neurons believed to be contained in 1 mm^3^ of mouse cortex. (right) The same as on the left, but instead of a pool of neurons of fixed size, each code is given a fixed total amount of energy. The energy is allocated to both passive maintenance of a neural population (with size equal to the population size of the code) and representation energy (increasing SNR). The shaded area is the same as on the left.

**Figure S2:**
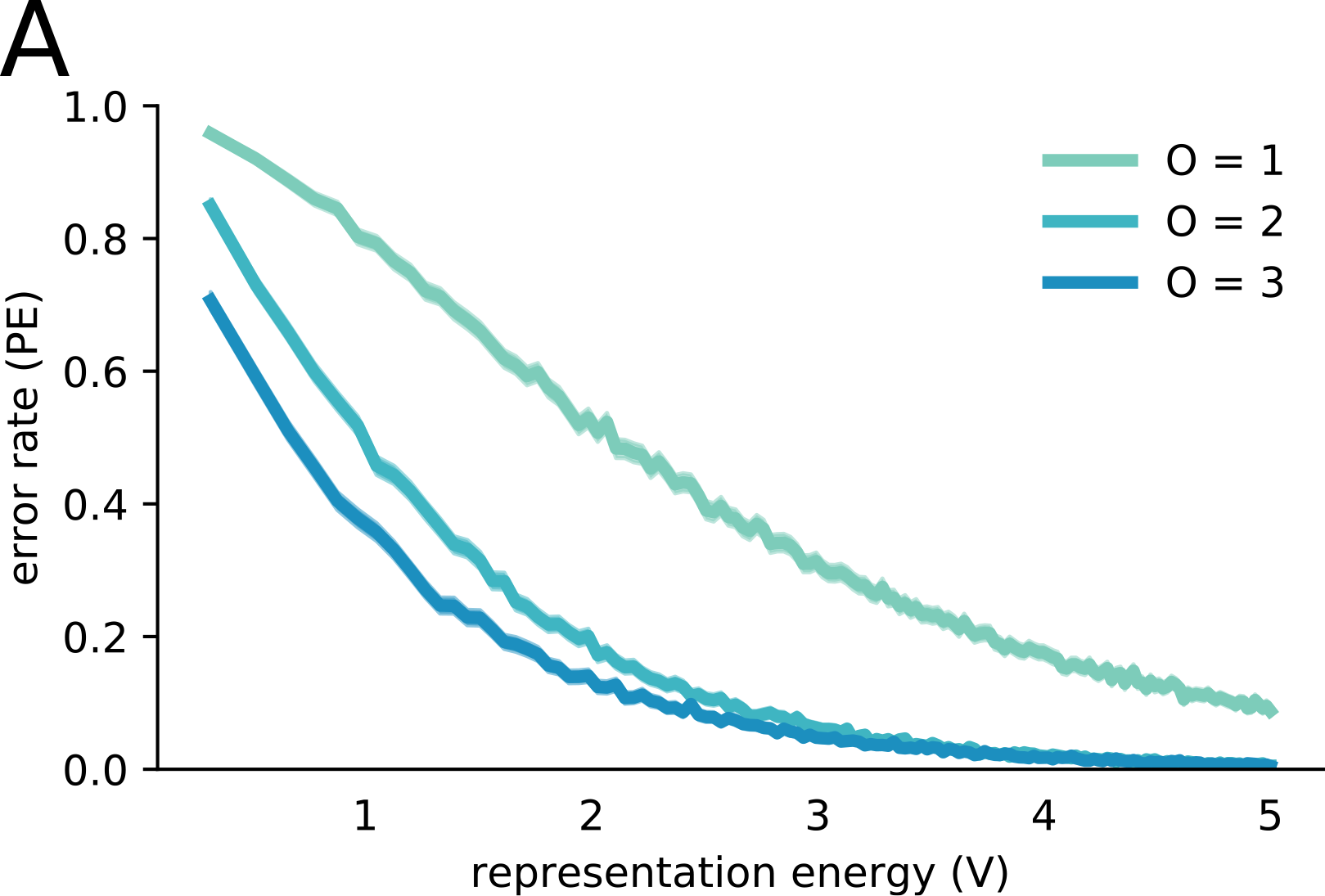
Channels with Poisson noise have similar performance to those with Gaussian noise, related to Figure 2. **A** The error rate (PE) as a function of representation energy (*V*) for codes with Poisson distributed noise, *K* = 3 and *n* = 5.

**Figure S3:**
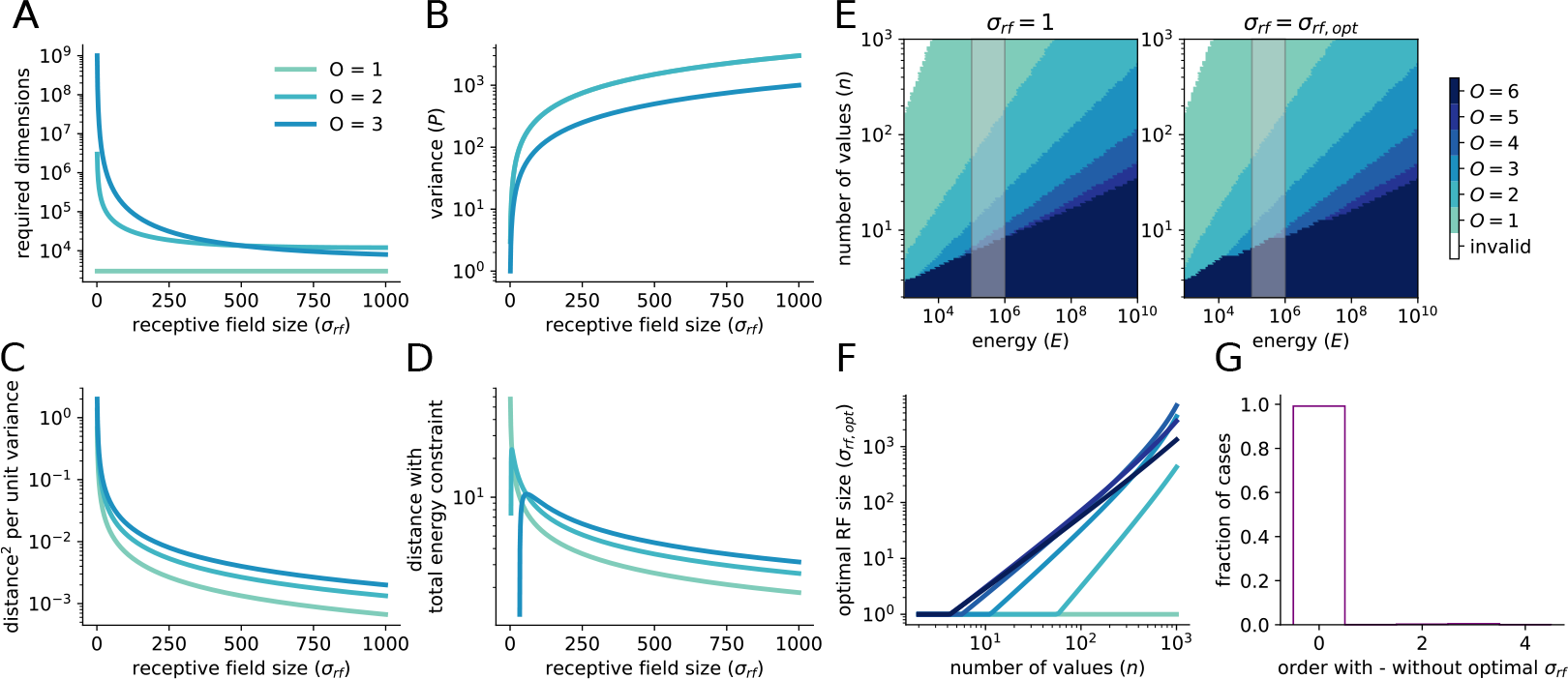
Changing response field (RF) size changes code properties, related to Figure 3. **a** The number of dimensions required to implement the code decreases by several orders of magnitude. **b** The power of the code increases by several orders of magnitude. **c** The tradeoff between minimum distance and code power remains constant if all codes are given the same RF size. **d** The RF size maximizing minimum distance under the total energy constraint differs between codes. **e** The code providing the highest minimum distance with *σ*_rf_ = 1 (left) and *σ*_rf_ = *σ*_rf,opt_ (right) as computed in Eq. S.2. They are only marginally different. **f** The optimal RF size for codes of different orders given features with different numbers of possible values. **g** Histogram of the differences in code order giving the highest distance from **e**.

**Figure S4:**
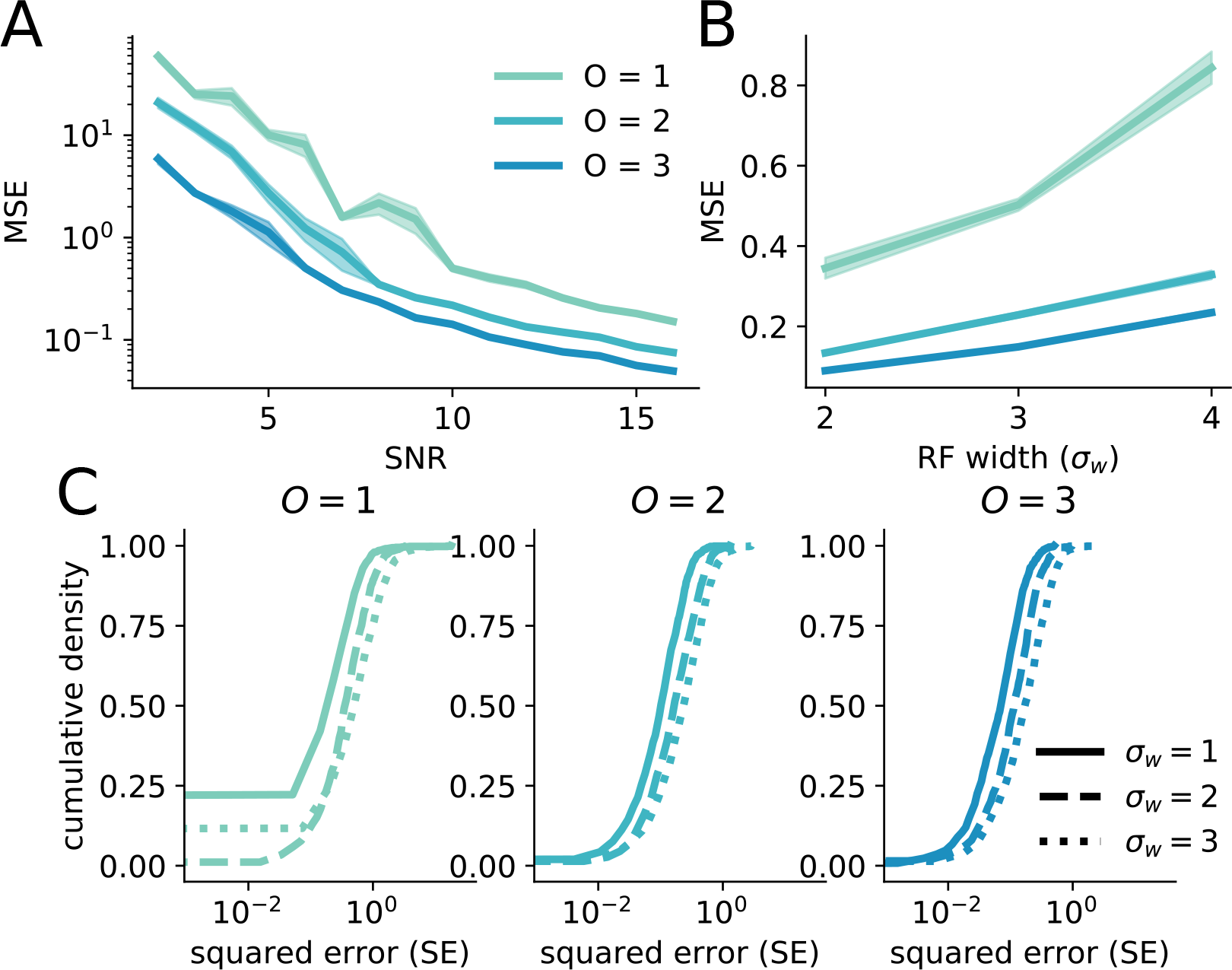
The benefits of mixed codes broadly generalize to continuous stimuli and RFs, related to Figure 3. **A** The MSE of codes of all orders with *K* = 3. The higher-order codes provide better performance than the lower-order codes. **B** MSE increases with RF size, which is contrary to the result in the discrete case (Figure 3d). **C** The cumulative distribution function of squared error for the three codes and for three different RF sizes.

